# A novel oral GyrB/ParE dual binding inhibitor effective against multidrug resistant *Neisseria gonorrhoeae* and other high-threat pathogens

**DOI:** 10.1101/2022.03.23.485573

**Authors:** Steven Park, Riccardo Russo, Landon Westfall, Riju Shrestha, Matthew Zimmerman, Veronique Dartois, Natalia Kurepina, Barry Kreiswirth, Eric Singleton, Shao-Gang Li, Nisha Mittal, Yong-Mo Ahn, Joseph Bilotta, Kristie L. Connolly, Ann E. Jerse, Joel S. Freundlich, David S. Perlin

## Abstract

Drug resistant *Neisseria gonorrhoeae* is a serious global health concern. New drugs are needed that can overcome existing drug resistance, as well as limit development of new resistance. We describe the small molecule tricyclic pyrimidoindole JSF-2414 [8-(6- fluoro-8-(methylamino)-2-((2-methylpyrimidin-5-yl)oxy)-9H-pyrimido[4,5-b]indol-4-yl)-2- oxa-8-azaspiro[4.5]decan-3-yl)methanol], which simultaneously binds to ATP binding regions of DNA gyrase (GyrB) and topoisomerase (ParE). JSF-2414 displays potent activity against *N. gonorrhoeae* including drug-resistant strains. A phosphate prodrug JSF-2659 was developed to facilitate oral dosing. In two different animal models of *Neisseria gonorrhoeae* vaginal infection, JSF-2659 was highly efficacious in reducing microbial burdens to the limit of detection. The parent molecule also showed potent *in vitro* activity against high-threat Gram positive organisms, and JSF-2659 was shown in a deep tissue model of VRSA and a model of *C. difficile*-induced colitis to be highly efficacious and protective. JSF-2659 is a novel drug candidate against high-threat multidrug resistant organisms with low potential to develop new resistance.

## BACKGROUND

Sexually transmitted infections due to *Neisseria gonorrhoeae* remain a significant global public health concern. Complications of gonorrhea affect women and men, and in women include pelvic inflammatory disease, ectopic pregnancy, and infertility, as well as increased transmission and acquisition of HIV ^1^. In 2012, the World Health Organization (WHO) estimated that there were 78 million cases among adults worldwide (https://www.paho.org/en/topics/sexually-transmitted-infections/gonorrhea.) In 2018, the U.S Centers for Disease Control and Prevention reported a total of 583,405 cases of gonorrhea with a national infection rate of 179.1 cases per 100,000 population, which reflects an increase of 63% since 2014 and the highest number since 1991 ^2^. For decades, gonorrhea was treated successfully using antimicrobials. Yet, there is now a high prevalence of *N. gonorrhoeae* strains that are resistant to common antimicrobial classes used for treatment including sulfonamides, penicillins, cephalosporins, tetracyclines, macrolides, and fluoroquinolones ^3^. Therapeutic failures with the extended-spectrum cephalosporins, such as cefixime and ceftriaxone, have created a major health crisis. In many countries, ceftriaxone is the only remaining empiric monotherapy for gonorrhea ^4, 5^. Given the high burden of gonococcal disease with resistance and the rapid emergence of resistant strains to monotherapy, antimicrobial therapy involving high dose ceftriaxone is recommended ^6^. Although, as anticipated, resistance to this regimen has also occurred ^7^. It is recognized that *N. gonorrhoeae* has evolved as a multidrug resistant superbug representing a major global public health concern ^7, 8^. The WHO has proposed a 90% reduction in gonorrhea globally ^9^, although achieving this goal will require overcoming the issue of antimicrobial resistance.

In recent years, new gonorrhea treatment regimens and drug candidates have been introduced to overcome and prevent resistance ^4^. The most promising class of new drug candidates interfere with DNA biosynthesis by inhibiting bacterial DNA gyrase (GyrB) and topoisomerase IV (ParE) via a unique mechanism ^4^. DNA gyrase and topoisomerase IV are closely related DNA topoisomerase type II enzymes that are essential for DNA synthesis. These enzymes function in tandem to catalyze topological changes in DNA during replication through supercoil unwinding and subsequent introduction of transient double-stranded DNA breaks and religation, and serve as the target for fluoroquinolone therapeutics ^10^. This dual targeting paradigm confers high susceptibility of new drug candidates against both fluoroquinolone-susceptible and resistant isolates, and further carries a very low probability for development of new resistance.

Zoliflodacin, a first-in class spiropyrimidinetrione ^11, 12^, and Gepotidacin, a triazaacenaphthylene inhibitor ^13^ leverage this dual-targeting approach and are novel, clinical-stage drug candidates that are completing phase 2 trials for the treatment of uncomplicated gonorrhea. These compounds inhibit bacterial DNA gyrase and topoisomerase IV by a novel mode of action that involves a binding site close to, but distinct from that of quinolones. Given their mechanism of action, these agents are broadly active against Gram-negative and Gram-positive bacteria including other biothreat agents such as *Streptococcus pneumoniae*, *Haemophilus influenzae*, *Clostridium perfringens*, and various *Shigella* species ^12–15^.

The tricyclic pyrimidoindoles are a new class of highly potent molecules that inhibit both GyrB and ParE (TriBE inhibitors) by binding at the highly conserved ATP-binding domain ^16^. As the ATP-binding region is separate and apart from the fluoroquinolone binding domain, it would not be subject to common resistance-associated target site mutations. The TriBE inhibitors demonstrate potent broad-spectrum Gram-negative and Gram-positive *in vitro* activity including against drug-resistant strains ^16^.

In this report, we describe the potent *in vitro and in vivo* properties of the tricyclic pyrimidoindole JSF-2414 and its oral pro-drug conjugate JSF-2659 against drug-sensitive and drug-resistant strains of *N. gonorrhoeae*, and drug-resistant Gram-positive pathogens including Methicillin and Vancomycin Resistant *Staphylococcus aureus* (MRSA & VRSA) and *Clostridioides difficile*.

## RESULTS

### Chemical and pharmacological properties of JSF-2414 and JSF-2659

The small molecule compound 8-(6-fluoro-8-(methylamino)-2-((2-methylpyrimidin-5-yl)oxy)-9H- pyrimido[4,5-b]indol-4-yl)-2-oxa-8-azaspiro[4.5]decan-3-yl)methanol (JSF-2414) (**Fig. 1**) was selected from a series of tricyclic pyrimidoindoles with potent activity (MIC <0.05 μg/ml) against the fastidious Gram-negative organism *Neisseria gonorrhoeae* and the Gram-positive organism *Staphylococcus aureus*. JSF-2414 has a molecular weight of 493.54 g/mol. Its solubility, metabolic stability, and *in vitro* intrinsic clearance (CLint) in mouse and human liver microsomes are summarized in **Fig. 1**. The *in vitro* metabolic stability of the compound in the presence of mouse or human liver microsomes was in an acceptable dosing range (i.e., t1/2 ≥ 60 min [PMID: 29311070]). The relatively low kinetic solubility of JSF-2414 (0.576 µM) was addressed by synthesizing a phosphate prodrug candidate JSF-2659 (**Fig. 1** and synthetic scheme in Supplement), which increased the solubility of the parent drug candidate 764-fold to 440 µM. The PK of JSF-2659 administered orally at 25 mg/kg showed a T1/2 = 2.03 ± 0.25 h and a Cmax ∼ 3500 ng/mL, while administration of JSF-2659 intramuscularly (IM) increased the T1/2 to 12.5 ± 0.25 h while the Cmax decreased slightly to ∼2400 ng/mL.

**Figure 1.**
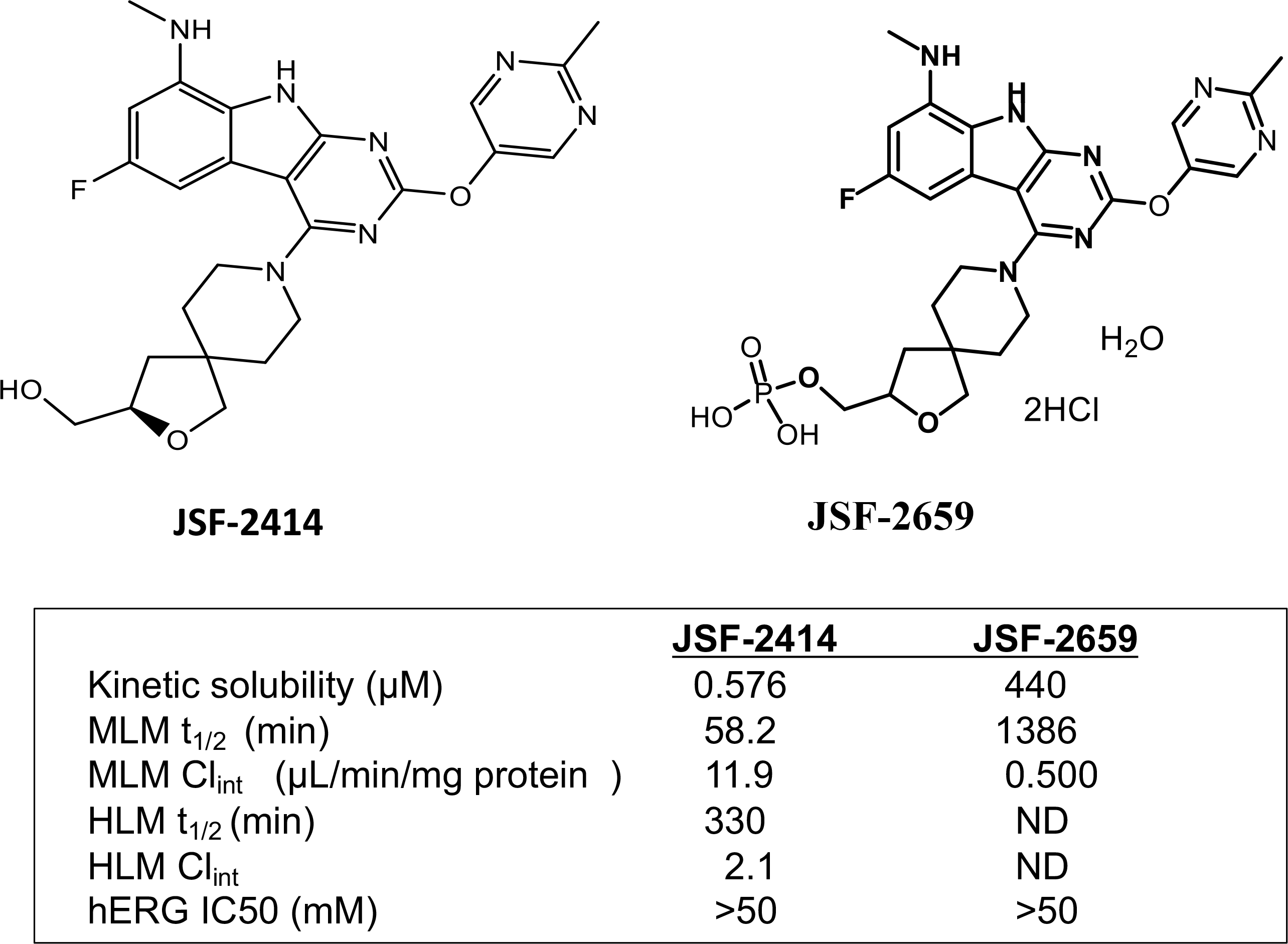
Preclinical development candidates. Chemical structures for JSF-2414 (8-(6- fluoro-8-(methylamino)-2-((2-methylpyrimidin-5-yl)oxy)-9H-pyrimido[4,5-b]indol-4-yl)-2- oxa-8-azaspiro[4.5]decan-3-yl)methanol and phosphate derivative JSF-2659 showing kinetic aqueous solubility and metabolic stability including half-life (t1/2) and intrinsic clearance (Clint) parameters with isolated mouse liver microsomes (MLM) and human liver microsomes (HLM).

### Antimicrobial activity against *Neisseria gonorrhoeae* and other high-threat drug- resistant pathogens

The antimicrobial properties of JSF-2414 (parent drug candidate) were evaluated by the Southern Research Institute against a collection of 96 *N. gonorrhoeae* clinical isolates including drug-resistant strains provided by the Centers for Disease Control and Prevention (CDC) under contract with the National Institutes of Health (See Methods.) An agar dilution method was used according to guidelines established by the Clinical and Laboratory Standards Institute, along with six control antibacterial agents (azithromycin, cefixime, ceftriaxone, ciprofloxacin, penicillin, and tetracycline) which served as reference controls. JSF-2414 was highly activity against all drug-susceptible and drug-resistant strains with an MIC90 = 0.006 µg/ml (range: 0.0005 –0.003 µg/ml.) Individual MIC values for each strain are listed in **Supplement Table 1**.

*In vitro* susceptibility testing was further performed against a highly diverse panel of Gram-positive and negative bacterial species. JSF-2414 showed prominent growth inhibition against 51 clinical MRSA strains with a modal MIC = 0.031 μg/ml (range 0.002–0.125 μg/ml). As JSF-2414 binds apart from the region of gyrase that confers resistance to fluoroquinolones, all ciprofloxacin-resistant strains with an MIC >4 μg/ml were fully sensitive (MIC = 0.043 μg/ml; range: 0.002 – 0.125 μg/ml; n=30). JSF-2414 also showed potent activity against VISA (MIC range 0.0156 – 0.156; n=8) and VRSA (MIC range 0.0156 – 0.063; n=7) strains (**Table 1**.)

**Table 1.** Susceptibility VRSA and VISA clinical isolates to JSF-2414

| Strains | JSF-2414<br>(µg/ml) | Vancomycin<br>(µg/ml) | Rifampicin<br>(µg/ml) |
| --- | --- | --- | --- |
| MRSA ATCC 43300 | 0.031 | 3.125 | <0.078 |
| **VRSA Strains** |  |  |  |
| VRSA 22522 | 0.031 | >200 | >10 |
| VRSA 22523 | 0.016 | 100 - 200 | <0.078 |
| VRSA 22524 | 0.016 - 0.031 | 200 | <0.078 |
| VRSA 22525 | 0.016 - 0.031 | 100 | <0.078 |
| VRSA 22526 | 0.0156 - 0.031 | <1.56 | <0.078 |
| VRSA 22527 | 0.031 | 1.56 - 3.13 | <0.078 |
| VRSA 22528 | 0.0156 - 0.031 | >200 | <0.078 |
| **VISA Strains** |  |  |  |
| VISA 21143 | 0.0156 - 0.031 | 12.5 | <0.078 |
| VISA 21156 | 0.063 | 3.13 - 6.25 | >10 |
| VISA 21157 | 0.031 - 0.063 | 12.5 | <0.078 |
| VISA 21161 | 0.0156 - 0.031 | 6.25 | <0.078 |
| VISA 21171 | 0.0156 - 0.031 | 3.125 | >10 |
| VISA 21178 | 0.0156 - 0.031 | 6.25 | <0.078 |
| VISA 21186 | 0.031 | 6.25 | >10 |
| VISA 21352 | 0.0156 - 0.031 | 6.25 | >10 |

JSF-2414 was also highly active against *Staphylococcus epidermidis* (MIC range 0.002–0.004 μg/ml; n=4), *Enterococcus faecium* (MIC range 0.004 – 0.008; n=3), vancomycin-resistant enterococci (VRE) (MIC range 0.002 – >0.5 μg/ml; n=8), and *E. faecalis* (MIC range 0.002 – 0.008; n=3), *B. anthracis* (MIC range 0.049 – 0.098 μg/ml; n=2). JSF-2414 was assessed for activity against a toxigenic *C. difficile* panel that included the common ribotypes 001, 002, 012, 014, 020, 038, 078, and 087 as well as the highly virulent ribotype 027. All *C. difficile* strains tested were highly susceptible to JSF-2414 with a MIC <0.125 μg/ml. In general, there was weak activity (MIC >0.5 μg/ml) against Gram negative pathogens *Escherichia coli, Pseudomonas aeruginosa, Klebsiella pneumoniae,* and *Acinetobacter baumannii* and select agents *F. tularensis and Y. pestis* (data not shown.)

In studies of *N. gonorrhoeae* and *S. aureus*, spontaneous resistance was not observed suggesting resistance frequencies below 5 x10^-9^.

### *Efficacy of* JSF-2659 in *Neisseria gonorrhoeae* vaginal colonization models

JSF- 2659 is rapidly and completely (>99%) converted by host phosphatases to its highly active form JSF-2414 following oral administration in mice (data not shown). The *in vivo* efficacy of JSF-2659 was initially assessed in a well-characterized estradiol-treated mouse model of cervico-vaginal infection at the Uniformed Services University (USU) ^17^ against multi- drug-resistant strain H041 ^18^, which carries resistance to extended-spectrum cephalosporins, tetracycline, macrolides, and several fluoroquinolones including ciprofloxacin, ^17^. The detailed experimental design of the model is show in **Supplement Fig. 1.** JSF-2659 administered at doses of 75 mg/kg TID (q6h), 250 mg/kg QD (once daily) or 250 mg/kg TID (q6h) showed a significant reduction in the percentage of mice infected colonized with strain H041 over 8 culture days after treatment (**Fig. 2 A,B**), as well as a reduction in tissue burden (CFU/ml recovered.) (**Fig. 2C**). The three dosing regimens showed a comparable reduction in the percentage of infected mice relative to the Gentamicin (GEN) positive control group. However, the average CFU/mL recovered was significantly reduced compared to the vehicle control only after 250 mg/kg TID treatment (**Fig. 2 B**). After Day 5 post-treatment, a significant reduction in bacterial burden was observed between all treatment groups and the vehicle control (**Fig 2 C**). All JSF- 2659 treatment groups had a comparable or greater reduction in bacterial burden relative to the untreated control or the GEN-treated control. It was observed that microbial burdens in some animals rebounded following treatment (**Fig. 2B**). However, in follow-up studies, there was no indication of resistance development or change in susceptibility (not shown).

**Figure 2.**
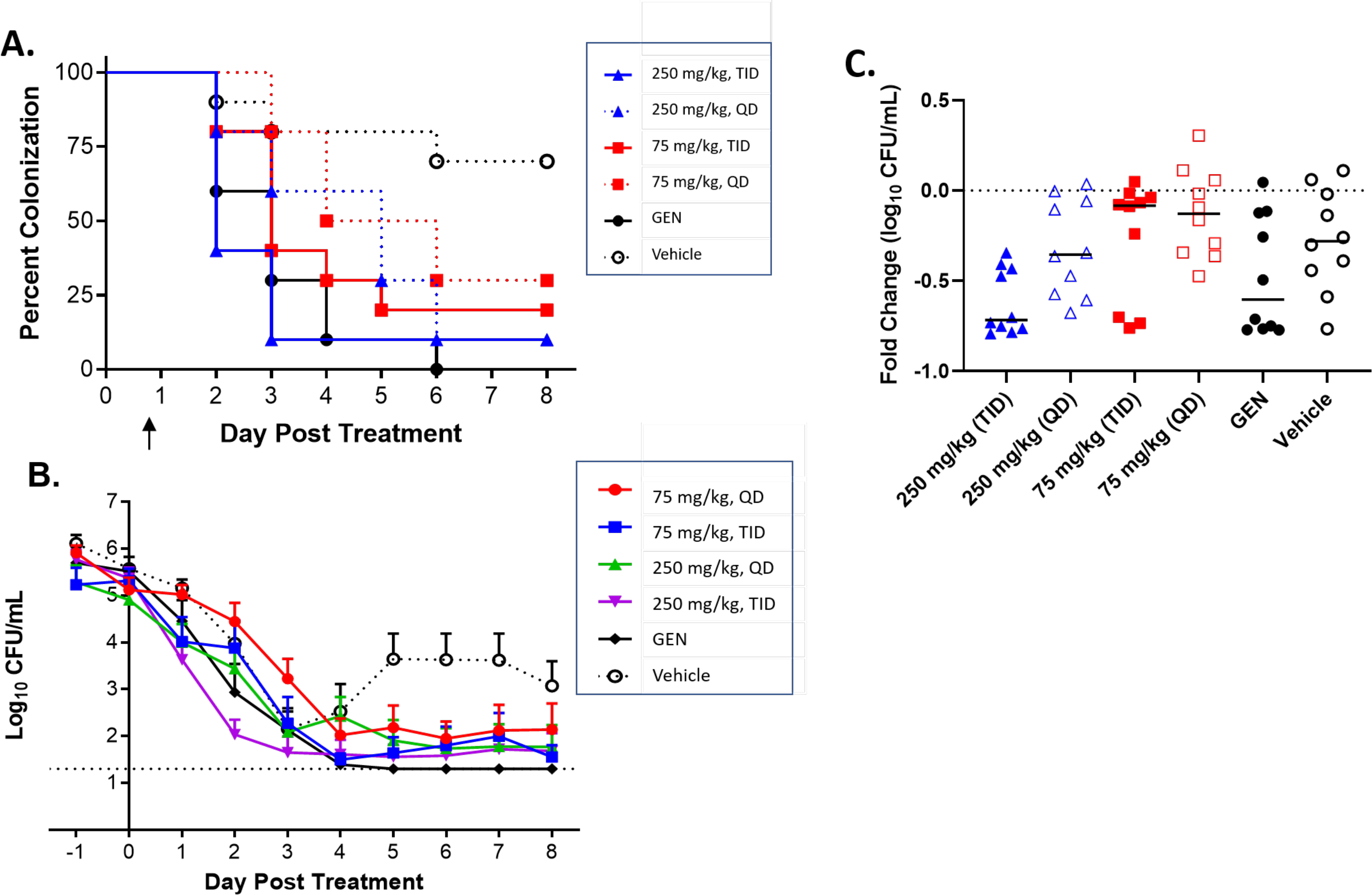
Dose-dependent efficacy of JSF-2659 against the multi-drug resistant *Neisseria gonorrhoeae* strain H041 in the murine vaginal infection model. Animals were vaginally inoculated with H041 bacteria and two days later, treated daily with JSF-2659 as shown in Suppl. Fig 1. Mice received daily oral doses of JSF-2659 at 75 mg/kg (blue) or 250 mg/kg (red), either TID (solid lines) or QD (dashed lines) over 8 days. Gentamycin at 48 mg/kg, 5 QD was used as a positive control; the control vehicle was used as the negative- control. **A.** A dose-dependent clearance rate was observed in JS-2659-treated mice as assessed by the percentage of culture-positive mice on eight consecutive days following treatment initiation. p values 0.02, 0.01, 0.003 and 0.0007 for the 250 mg/kg, QD, 75 mg/kg, TID, 75 mg/kg QD and gentamycin control group versus the vehicle control group, respectively (Log-rank (Mantel-Cox test). **B.** The average number of CFU recovered per mL of vaginal swab suspension declined over time in all antibiotic treatment groups compared to the vehicle control group, with the differences between the 250 mg/kg, TID group and gentamycin control groups significantly different than the vehicle control group (p 0.004 and 0.007, respectively; 2-way ANOVA with repeated measures, Bonferroni post-hoc analysis); **C.** Fold change in CFU/mL between Day 0 (pre-treatment) and 48h post-treatment showed a trend towards a significant reduction for the 250 mg/kg, TID treatment group (p = 0.058).

The promising efficacy of JSF-2659 was independently assessed via PO and intramuscular administration (IM) in a different *Neisseria gonorrhoeae* vaginal colonization model using the antibiotic-susceptible strain ATCC 700825. In this model, ovariectomized female BALB/c mice were used and mice were given subcutaneous (SC) injections of 17β-estradiol. The influence of vaginal bacteria was minimized by treating animals BID 2 days prior to infection with streptomycin and vancomycin, along with trimethoprim sulfate at 0.4 mg/mL (see Methods). PO dosing and an alternative treatment route IM were tested with JSF-2659 at 75 and 250 mg/kg QD at 2 h after vaginal inoculation with bacteria. A single dose IM route was evaluated because of the long PK (T1/2 = 12.5 h) observed for JSF-2659 following IM administration. At 26 h post infection and 2 h post-therapy, a significant dose-dependent reduction in the bacterial counts was observed resulting in a 3.51 log10 killing effect to the limit of detection (LOD) following PO administrations of JSF-2659 at 75 and 250 mg/kg TID and JSF-2659 at 250 mg/kg QD, as well as 3.51 and 3.39 log10 killing effect following the IM administrations of JSF-2659 at 75 and 250 mg/kg QD, respectively, relative to control (*p*<0.05). A 1 log10 killing effect was observed with the PO administration of JSF-2659 at 75 mg/kg QD relative to the baseline group (*p* < 0.05). A 3.17 log10 killing effect was observed with the PO administration of reference ciprofloxacin (CIP) at 12.5 mg/kg QD relative to the baseline group (*p* < 0.05). No significant reduction in the bacterial counts was observed with the IP administration of GEN at 48 mg/kg QD relative to the vehicle control group or the baseline group at 26 h (**Fig. 3 A,B**). Similar reductions in bacterial burdens were observed at 72 h (**Fig. 3 C,D**) and 170 h (**Fig. 3 E,F**) following initiation of treatment. In summary, >3 log10 killing effects relative to the baseline group were observed in animals sacrificed at 26, 76 and 170 h after infection with (1) PO administrations of JSF-2659 at 75 and 250 mg/kg TID; (2) PO administrations of JSF-2659 at 250 mg/kg QD; (3) IM administrations of JSF2659 at 75 and 250 mg/kg QD; and (4) PO administration of reference CIP at 12.5 mg/kg QD.

**Figure 3.**
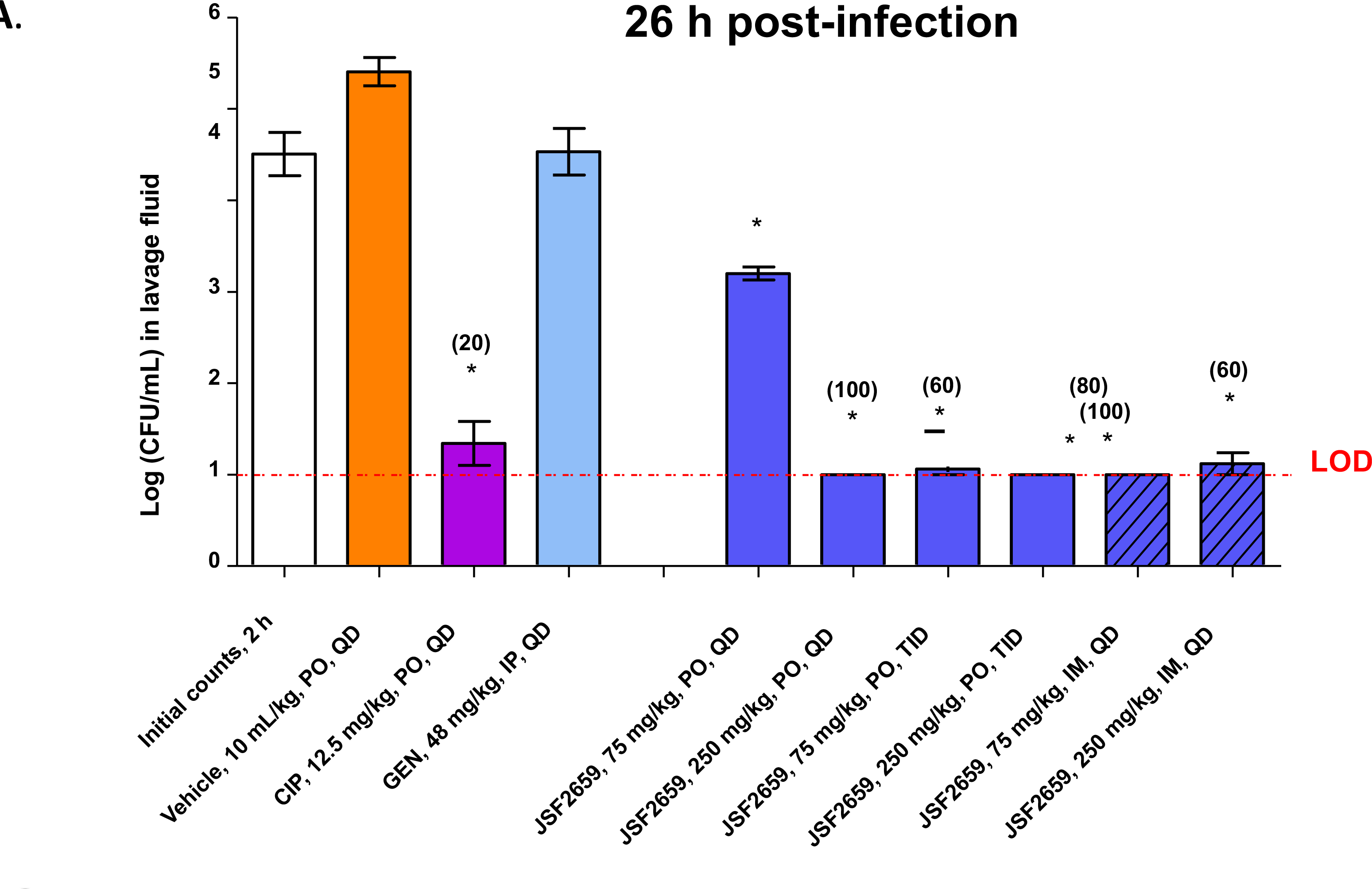
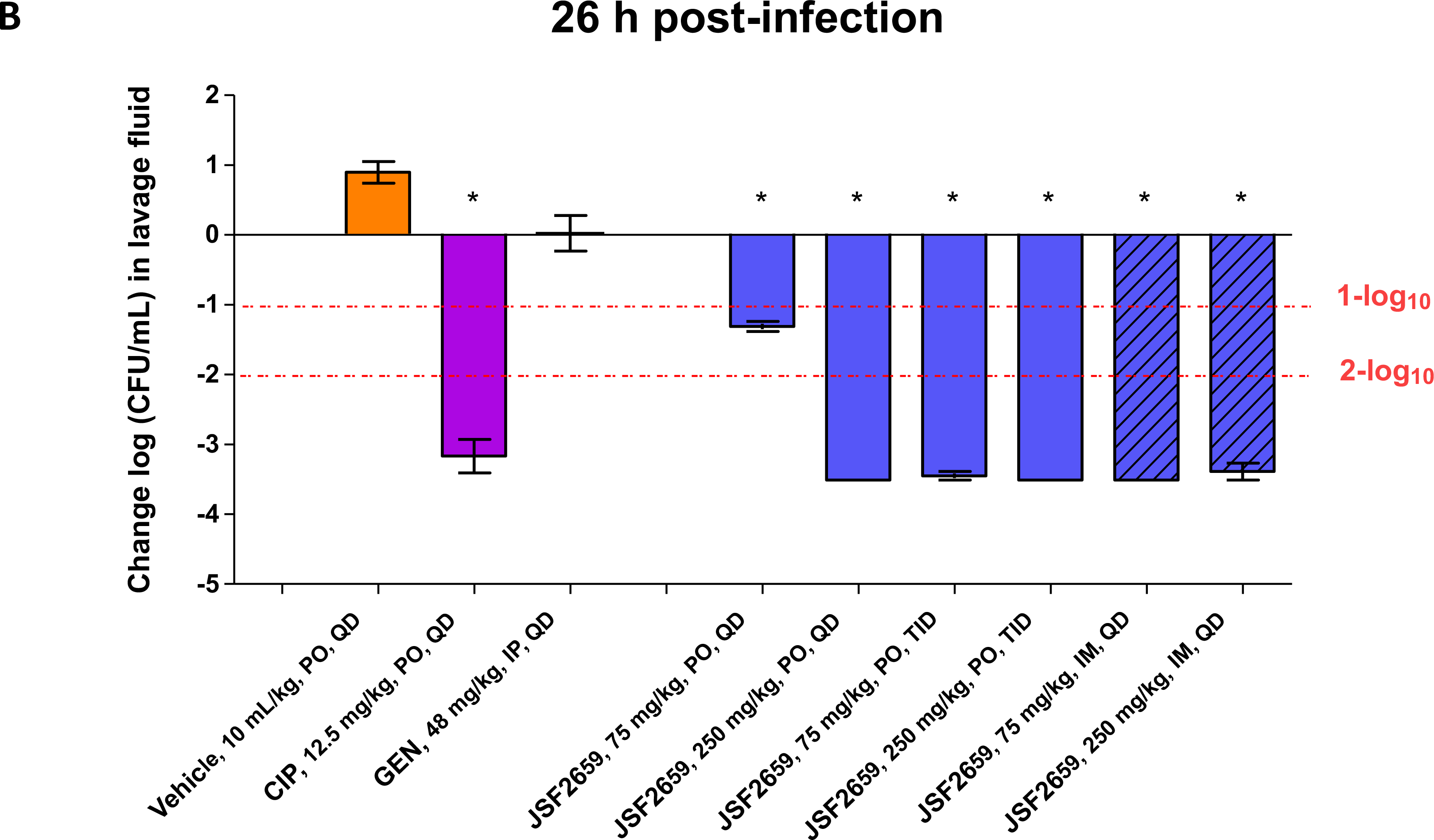
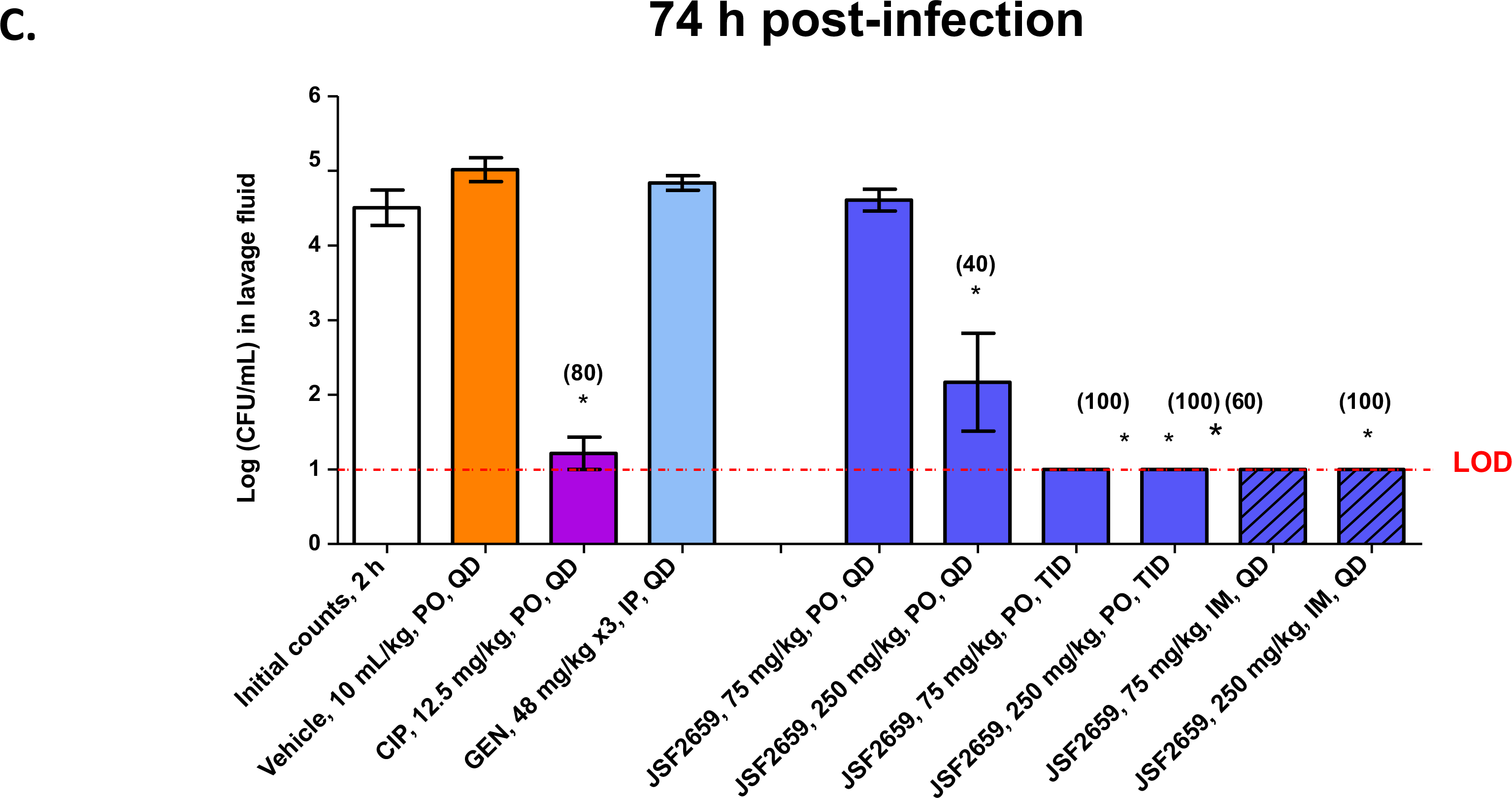
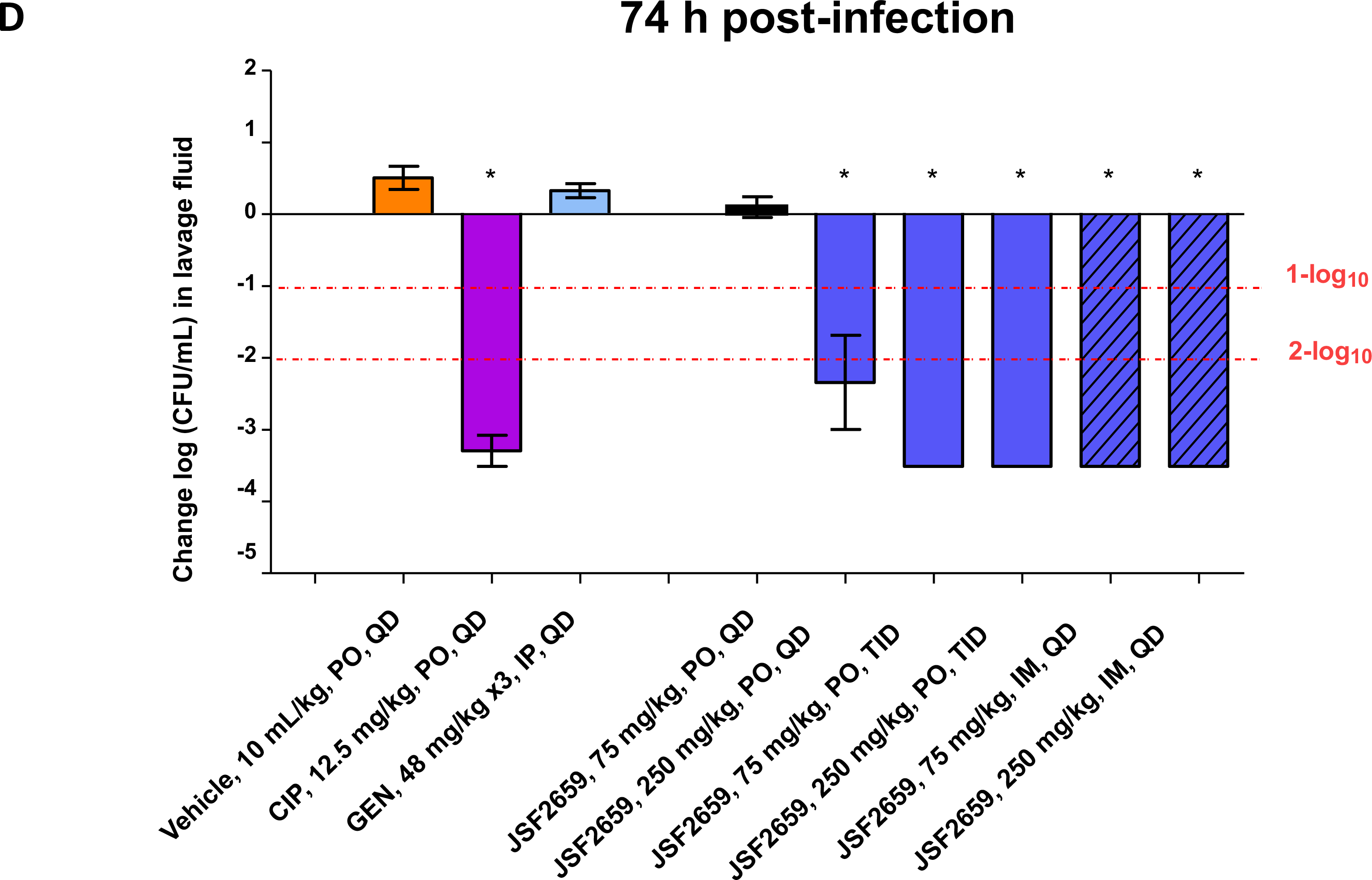
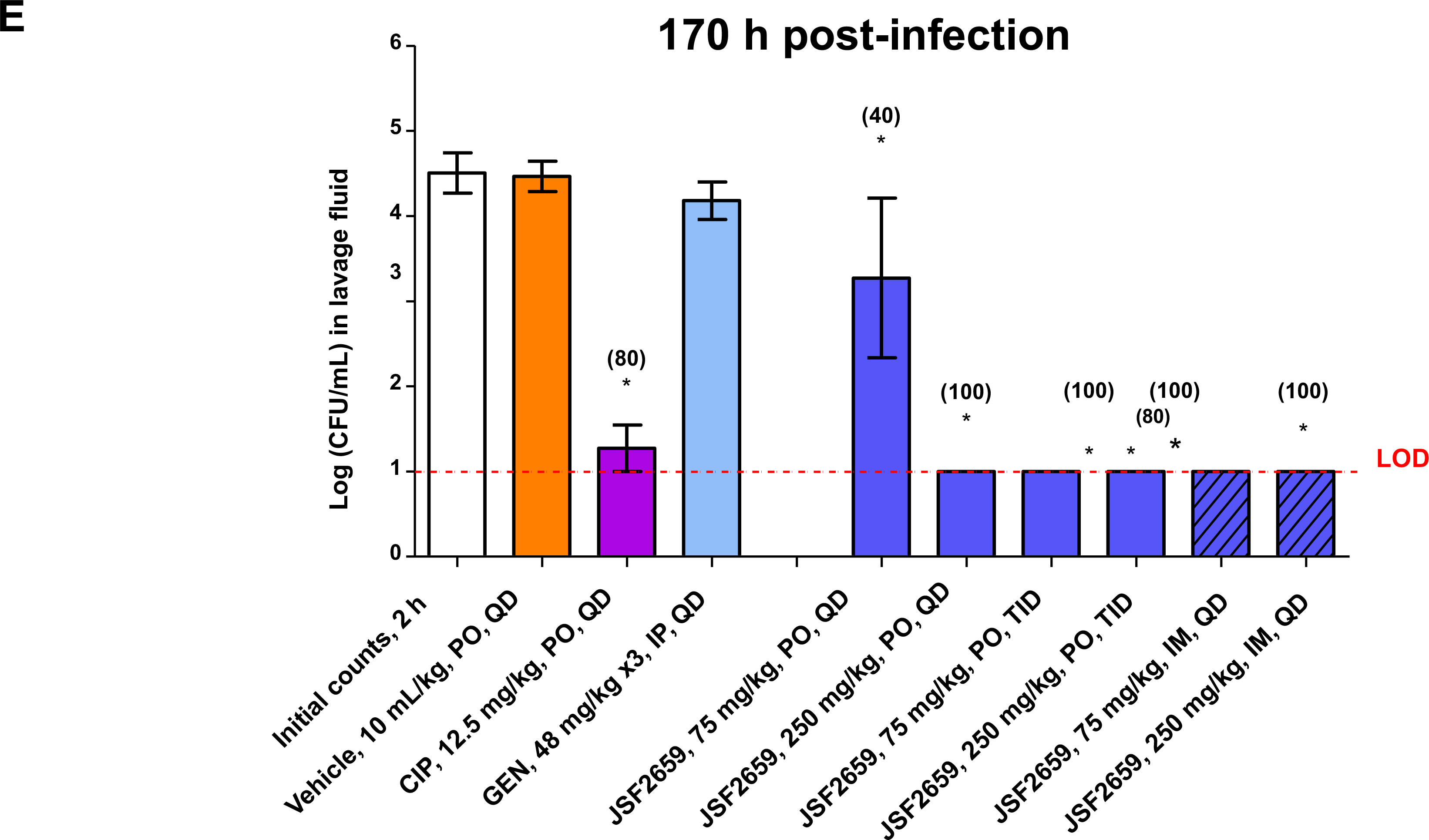
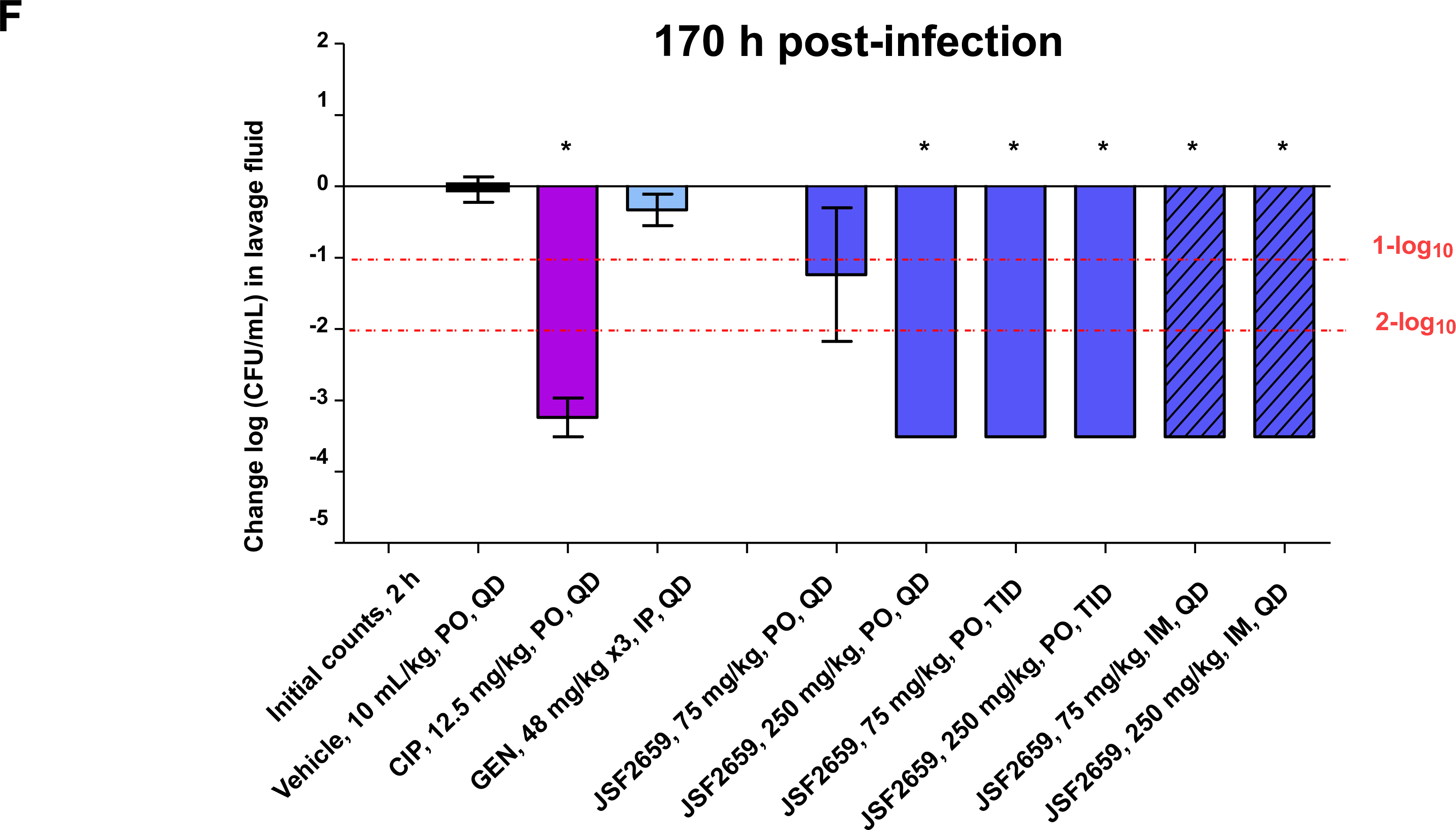
Effects of JSF2659, ciprofloxacin and gentamicin in a *N. gonorrhoeae* (ATCC 700825) intravaginal infection model using ovariectomized BALB/c mice. Total bacterial counts (**A,C,E**) and a net change in counts (**B,D,F**) is shown for vaginal lavage fluids following test article treatment relative to the initial 2 h counts at the time of dosing. JSF2659 at 75 and 250 mg/kg, was administered orally (PO) with two dosing schedules, including once (QD) at 2 h after infection, and three times (TID) with an 6 h interval at 2, 8, and 14 h after infection. In addition, JSF2659 at 75 and 250 mg/kg was administered intramuscularly (IM) once (QD) at 2 h after infection. Two reference compounds were applied in this study, ciprofloxacin (CIP) and gentamicin (GEN). Ciprofloxacin was administered orally (PO) at 12.5 mg/kg once (QD) at 2 h after infection. Gentamicin was administered intraperitoneally (IP) at 48 mg/kg once (QD) at 2 h after infection for seven consecutive days. Animals were sacrificed at 26 h (**A,B**), 72 (**C,D**) and 170 h (**E,F**) after infection. Vaginal lavage was performed and the bacterial suspensions were plated onto chocolate agar to determine the *N. gonorrhoeae* counts. (*): Significant difference (*p* < 0.05) compared to the respective vehicle control was determined by one- way ANOVA followed by Dunnett’s test.

### Efficacy of JSF-2659 against VRSA in a deep soft tissue infection model

JSF-2414 displayed potent *in vitro* activity against strains of MRSA, VRSA and VISA (**Table 1**). The oral efficacy of JSF-2659 against Vancomycin Resistant *Staphylococcus aureus* (VRSA) was assessed in a neutropenic murine thigh infection model. Female ICR mice were rendered neutropenic with cyclophosphamide treatment and then inoculated intramuscularly with VRSA (VRS-2) at 1.03 x 10^5^ CFU/mouse. JSF-2659 at 100 and 250 mg/kg was administered orally (PO) with two dosing schedules, including QD at 2 h after infection, and TID with a 6 h interval at 2, 8, and 14 h after infection. Linezolid and vancomycin served as reference standards. JSF-2659 PO administrations at 250 mg/kg QD and 100 and 250 mg/kg, resulted in significant bacterial count reductions 0.95 and 4.42 log10, respectively, relative to the vehicle control group (**Fig. 4**). When administered PO TID at 100 and 250 mg/kg, JSF-2659 yielded further bacterial burden reductions of 4.84 and 6.26 log10, respectively (**Fig. 4B**). The PO and SC administrations of respective reference agents, linezolid at 50 mg/kg BID and vancomycin at 100 mg/kg TID, yielded a reduction in bacterial counts of 3.41 and 2.09 log10, respectively.

**Figure 4.**
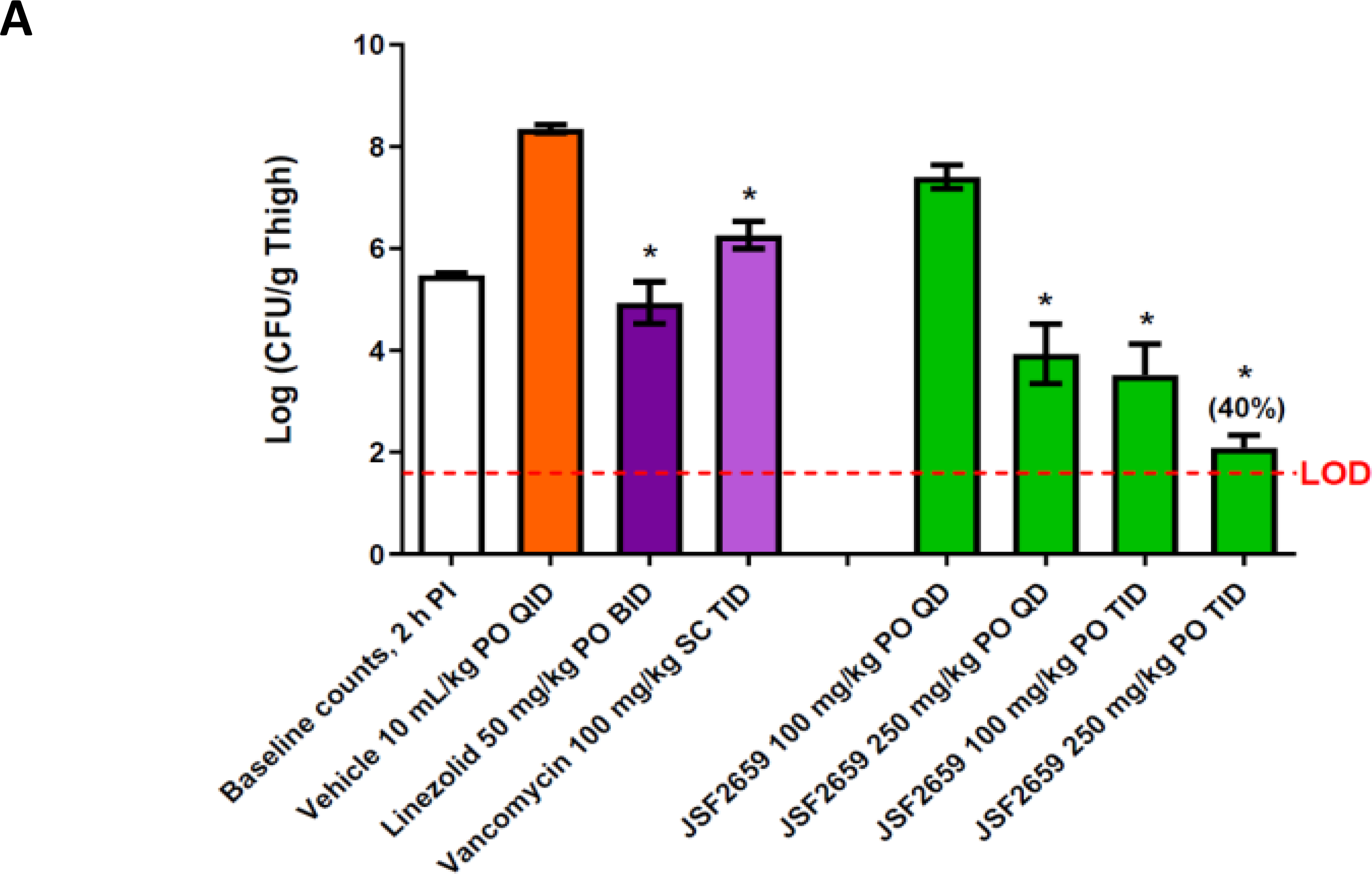
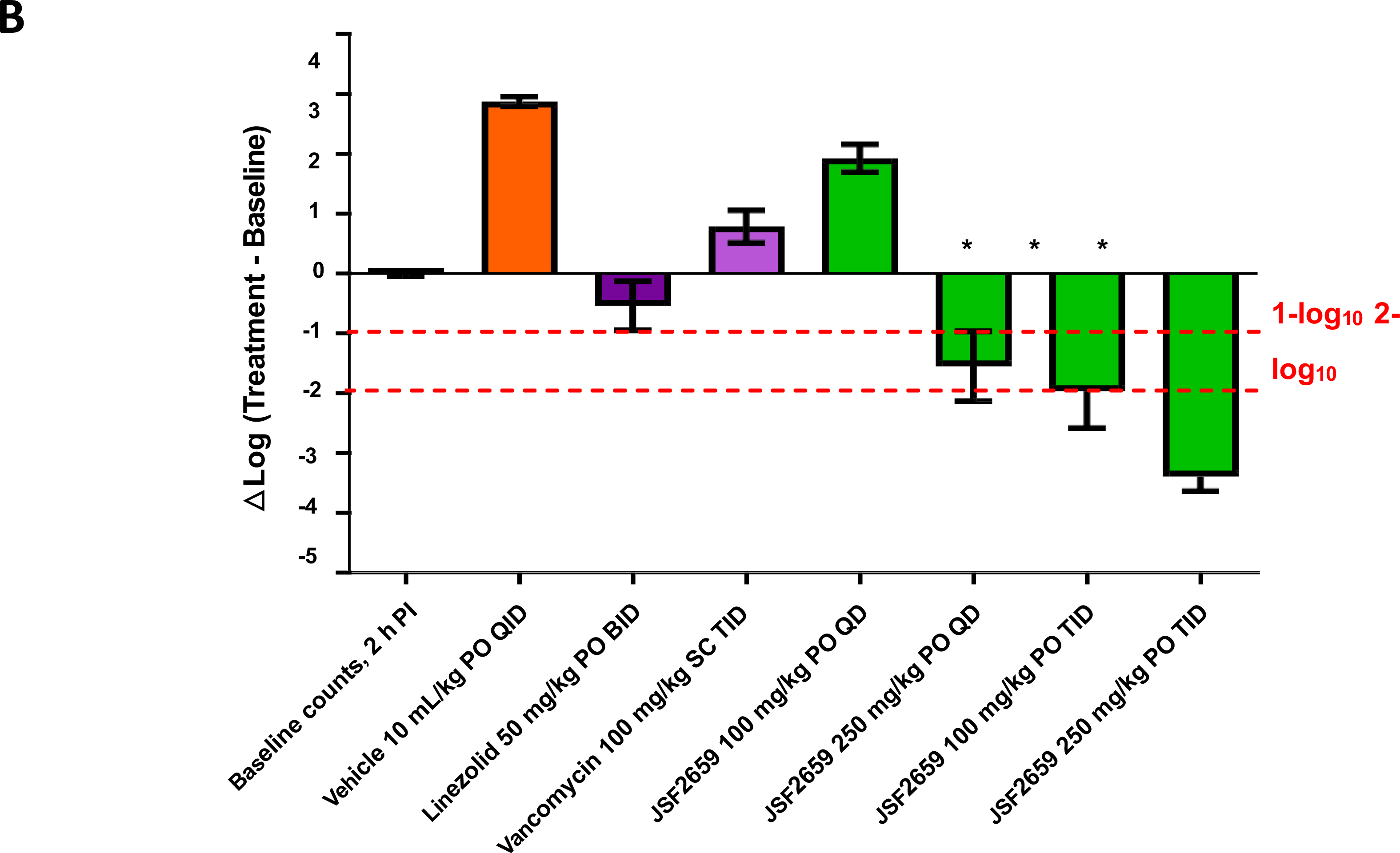
Effects of JSF2659, linezolid and vancomycin in the *S. aureus* (VRSA, VRS-2) thigh infection model with neutropenic mice. Mice were induced neutropenic with cyclophosphamide treatment and were then inoculated intramuscularly with an inoculation density of 1.0 x 10^5^ CFU/mouse. JSF-2659 at 100 and 250 mg/kg, was administered orally (PO) with two dosing schedules, including once (QD) at 2 h after infection, and three times (TID) with an 6 h interval at 2, 8, and 14 h after infection. Two reference compounds were applied in this study, linezolid and vancomycin. Linezolid was administered orally (PO) at 50 mg/kg twice (BID) at 2 and 14 h after infection. Vancomycin was administered subcutaneously (SC) at 100 mg/kg three times (TID) at 2, 8 and 14 h after infection. Animals were sacrificed at 2 or 26 h and the thigh tissues were harvested and the total bacterial counts (CFU/g)(**A**) and change in counts (**B**) in thigh tissues were compared. (*) Significant difference (*p* < 0.05) compared to the vehicle control was determined by one- way ANOVA followed by Dunnett’s test. Linezolid was administered orally (PO) at 50 mg/kg twice (BID) at 2 and 14 h after infection and Vancomycin was administered subcutaneously (SC) at 100 mg/kg TID at 2, 8 and 14 h after infection served as reference controls. One infected but non-treated group was sacrificed at 2 h after infection for the initial bacterial counts. Mice from the treatment of test article, reference compounds, and vehicle control groups were sacrificed at 26 h after infection. The thigh tissue was excised for bacterial enumeration, CFU/gram. One-way ANOVA followed by Dunnett’s comparison test was performed to assess statistical significance (*p* < 0.05) in the bacterial counts of the treated animals compared to the vehicle control group.

Efficacy of JSF-2659 against *C. difficile* in a colitis model

The protective efficacy of JSF-2659 in a hamster *C. difficile* colitis model was evaluated. The study was performed with a lethal dose (LD90-100) of strain *C. difficile* BAA-1805 (ribotype 027/NAP1/BI.) Test animals were pretreated with a single subcutaneous (SC) administration of clindamycin at 10 mg/kg on Day -1 to render the animals vulnerable to *C. difficile* infection. On Day 0, animals were inoculated orally with *C. difficile* BAA-1805 at 5.8 x 10^5^ spores/animal. JSF- 2659 at 20, 100, and 250 mg/kg, and reference agent, vancomycin at 20 mg/kg, were administered PO twice daily (BID) with an 8 h interval starting at 16 h after infection for five consecutive days. Animal mortality and body weight changes were monitored for 14 days. Infection with *C. difficile* resulted in 100% mortality during the 14-day observation period (**Fig. 5A**). All test animals survived the complete study period after treatment with JSF-2659 at 20, 100, and 250 mg/kg PO BID for five days (100% survival for all groups, *p* <0.05) (**Fig. 5A**). The PO administration of vancomycin at 20 mg/kg resulted in 100% survival over the treatment period (*p* < 0.05). A loss of significant body weight for test animals relative to positive control was not observed during the 14-day observation period and was consistent with compound efficacy (**Fig. 5B**).

**Figure 5.**
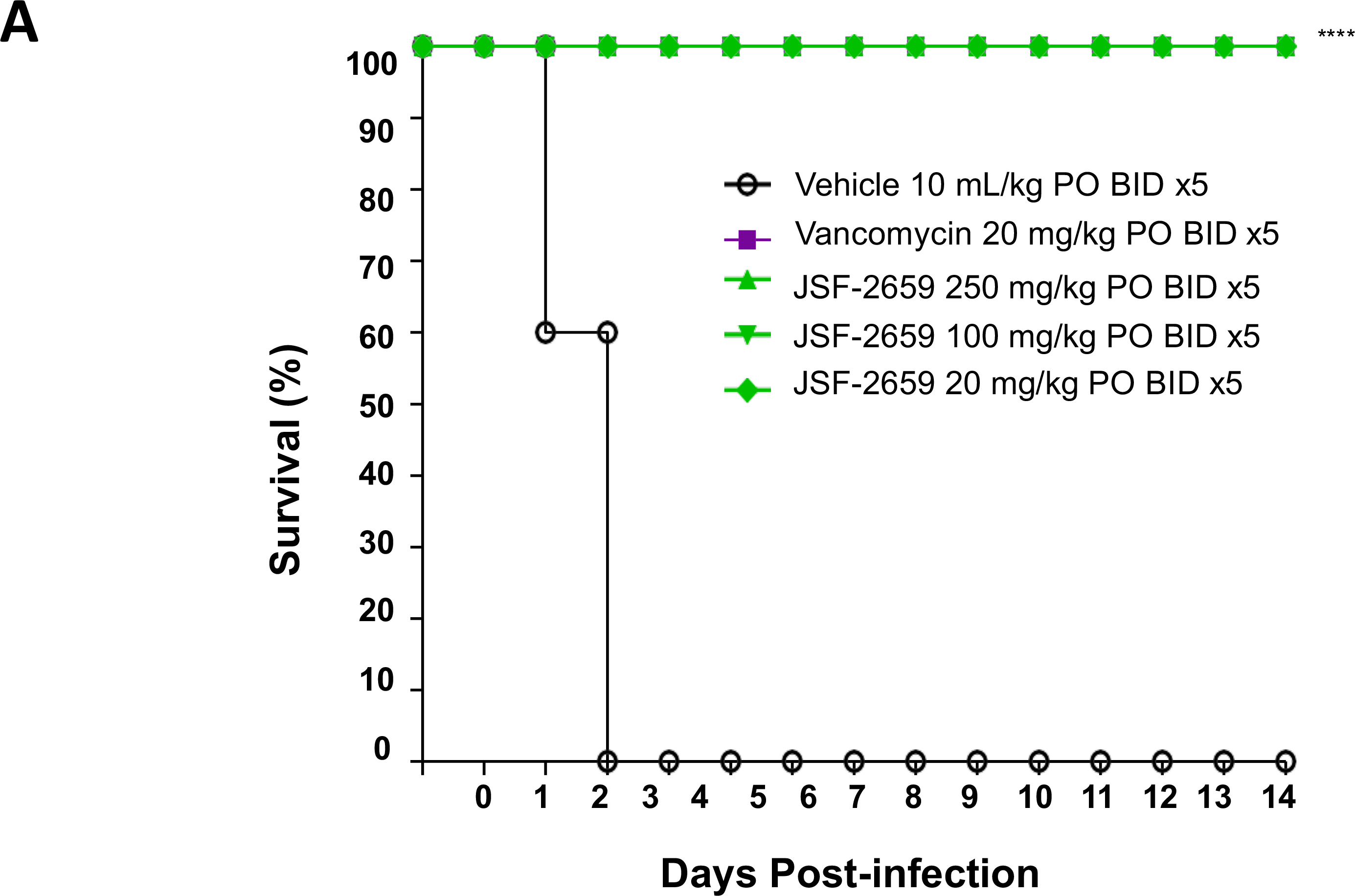
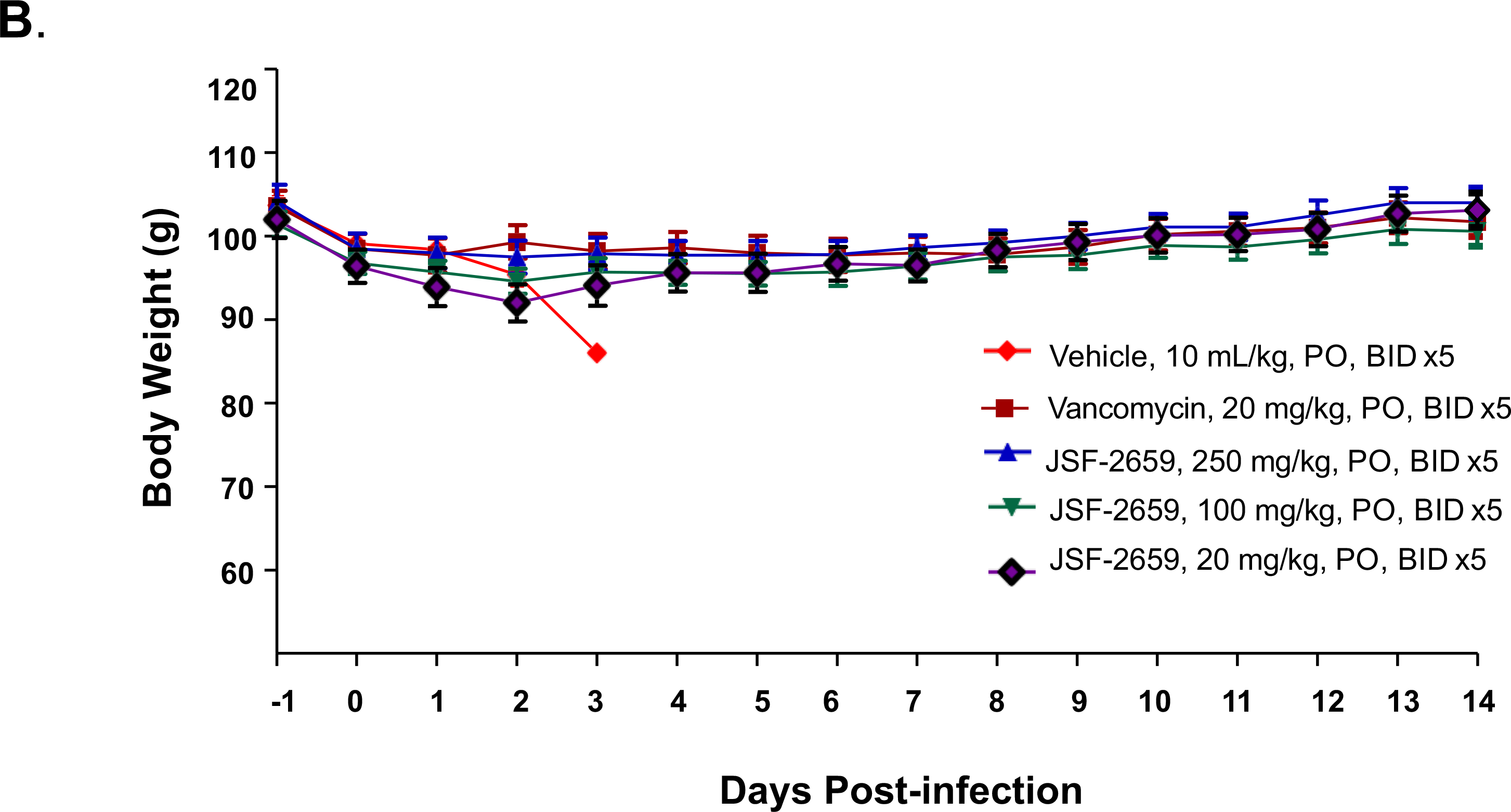
Kaplan-Meier plot of 14-day survival in the *C. difficile* ATCC BAA-1805 hamster colitis model. Animals were inoculated orally (PO) with *C. difficile* ATCC BAA-1805 at 5.8 × 10^5^ spores/animal. Test article JSF2659 at 20, 100, and 250 mg/kg, and reference agent, vancomycin at 20 mg/kg, were administered orally (PO) twice daily (BID) with an 8 h interval starting at 16 h after infection for five consecutive days. Animal mortality was monitored for 14 days (**A**). *: Indicated a significant increase (*p* < 0.05) in the survival rates of the treated animals compared to the vehicle control group at day 14 time point as determined by Fisher’s exact test. Test article, JSF2659 at 20, 100, and 250 mg/kg, and reference agent, vancomycin at 20 mg/kg, all showed significance and are indicated. **B.** Body weight changes of *C. difficile* ATCC BAA-1805 infected hamsters. Animals were inoculated orally (PO) with *C. difficile* ATCC BAA-1805 at 5.8 × 10^5^ spores/animal. Test article, JSF2659 at 20, 100, and 250 mg/kg, and reference agent, vancomycin at 20 mg/kg, were administered orally (PO) twice daily (BID) with an 8 h interval starting at 16 h after infection for five consecutive days. Body weight changes were monitored for 14 days.

## DISCUSSION

Despite the high worldwide burden of gonorrhea infections, the number of effective treatment options for gonorrhea infections is limited and further diminished by the emergence of multidrug resistance. This has prompted the WHO to place *N. gonorrhoeae* on a global priority list of antibiotic-resistant pathogens for which there is an urgent need to develop novel antimicrobial therapeutics ^19^. In this report, we describe the potent *in vitro* and *in vivo* antimicrobial properties of JSF-2414, a novel tricyclic pyrimidoindole that binds to the ATP binding domains of both DNA gyrase (GyrE) and topoisomerase IV (ParE). This tricyclic pyrimidoindole (**Fig. 1**) emerged from a study to identify a low molecular weight fragment scaffold with suitable hydrogen-bond donor/acceptor moieties that could engage the ATP adenine-binding aspartate and structural water in the active site pocket of *Enterococcus faecalis* GyrB ^16^. As gyrase and topoisomerase are well-established pharmacologic targets, targeting the ATP binding site is important because it is separate from the drug binding site of fluoroquinolones and resistance mediated mutations within the site ^20, 21^. It is not surprising that JSF-2414 was highly active, MIC range: 0.0005 – 0.003 µg/ml against a CDC panel (n=100) of clinical isolates of *N. gonorrhoeae* displaying a wide variety of drug resistance including fluoroquinolone resistance (Supplement, Table 1). This potent level of activity likely reflects the nature of its target and binding properties, which were originally optimized from structure-based studies ^16^.

The development of drugs targeting the ATP-binding domain is not new, as discovery programs for decades have focused on inhibitors (e.g., benzothiazole, tetrahydrobenzothiazole, etc.) that target the ATP-binding/hydrolysis sites on GyrB and ParE ^22–27^. Indeed, in the 1950s, the natural product novobiocin was reported as potently bactericidal to Gram-positive bacteria via inhibition of the ATP-binding domain of GyrB. It was licensed for clinical use under the tradename Albamycin (Pharmacia and Upjohn) in the 1960s but was withdrawn due to poor efficacy. Furthermore, its weak binding to ParE resulted in enhanced development of resistance. Newer efforts have also not been successful in generating an inhibitor series with broad spectrum antibacterial activity or advancing a molecule into the clinic. Most recently, the tetrahydroquinazoline and 4,5,6,7- tetrahydrobenzo[1,2-d] thiazole scaffolds were originally identified as low micromolar inhibitors of the DNA gyrase ATP-binding domain ^25^. But the compounds showed modest antibacterial activity, appeared to be substrates for efflux transporters and poorly penetrated the cell wall ^25^. In contrast to these efforts, JSF-2414, the tricyclic pyrimidoindole in this study, was highly active against a broad range of clinical isolates displaying a variety of resistance mechanisms (**Supplement Table 1**).

Potent but more modest MICs have also been reported for later stage clinical candidates that target more classical regions of gyrase. The spiropyrimidinetrione zoliflodacin, based on mapping of resistance associated mutations in GyrB, is presumed to bind to the same pocket in gyrase as fluoroquinolones ^12^. However, it inhibits gyrase through a separate mechanism ^28^, which enables it to be effective against most fluroquinolone resistant strains ^12^. In a multi-laboratory quality study of zoliflodacin against standardized ATCC strains by eight independent laboratories using CLSI document M23, the agar dilution MIC QC range for zoliflodacin against the *N. gonorrhoeae* QC strain ATCC 49226 was defined as 0.06 – 0.5 μg/ml ^29^. In a separate study, zoliflodacin showed potent antibacterial activity against multi-drug-resistant strains of *N. gonorrhoeae* with MICs ranging from ≤0.002 – 0.25 μg/mL ^12^. The triazaacenaphthylene inhibitor Gepotidacin has been shown to form an antibacterial complex with *S. aureus* DNA gyrase and DNA, demonstrating a novel mechanism of inhibition that overcomes fluoroquinolone resistance. The inhibitor was demonstrated to bind close to the active site for fluoroquinolone binding where it appears to span DNA and a transient non-catalytic pocket on the GyrA dimer ^30^. A multi-laboratory quality assurance study demonstrated a modal MIC of 0.5 μg/ml ^31^.

A hallmark of DNA gyrase inhibitors is their broad-spectrum activity especially against Gram-positive organisms. JSF-2414 showed potent cross-activity against MRSA with an MIC = 0.031 mg/ml (range 0.002 – 0.125 μg/ml) including VISA and VRSA strains (Table 1) with comparable activity against 30 clinical isolates with fluoroquinolone resistance MIC>4 µg/ml. It showed comparable high activity (low MICs) against *S. epidermidis*, *E. faecium*, VRE, *E. faecalis*, *B. anthracis,* MDR*-*TB, and *C. difficile*. Gepotidacin showed *in vitro* activity with an MIC90 of 0.5 μg/ml and equivalent activity against *S. pneumoniae* ^13^. Similarly, zoliflodacin in studies involving thousands of clinical isolates showed MIC values of 0.12 - 0.5 μg/ml for *S. aureus* ATCC 29213, 0.25 - 2 µg/ml for *E. faecalis* ATCC 29212, 0.12 – 0.5 μg/ml for S*. pneumoniae*, and 0.12 – 1 μg/ml for *Haemophilus influenzae* ^29, 32^. For all of the inhibitors, activity was less against prominent Gram-negative organisms like *E. coli* and *Pseudomonas aeruginosa.* For JSF-2414, gepotidacin and zoliflodacin, MIC values were 0.5, 4 and 4 µg/ml versus *E. coli*, respectively ^13, 29^.

To investigate the *in vivo* potential of JSF-2414 against *N. gonorrhoeae* infection, a phosphate prodrug candidate JSF-2659 (**Fig. 1**) was developed (Supplement) for oral dosing and *in vivo* efficacy of JSF-24214 was demonstrated in two different *N. gonorrhoeae* genital tract infection models. In one model, dosing of JSF-2659 at 75 mg/kg TID (q6h), 250 mg/kg QD or 250 mg/kg TID (q6h) showed a significant reduction in the percentage of mice infected with MDR strain H041 over 8 culture days after treatment (*p* ≤ 0.04 for 75 TID, 250 TID.) (**Fig. 2).** In another model dose-dependent oral administration of JSF-2659 profoundly reduced the burden of *N. gonorrhoeae* (ATCC 700825) by ∼4.5 log10 to the limit of detection (LOD.) Again, a PO dose-dependent response was observed with maximal reduction at 250 mg/kg QD, 75 mg/kg TID or 250 mg/kg TID at 26, 74 or 170 h post-administration (**Fig. 3**). A single IM administration at 75 or 250 mg/kg was effective in eliminating viable burdens (LOD) (*p* < 0.05 compared to vehicle control.)These promising data suggest that high exposures of the drug candidate are critical for the strong pharmacodynamic response. This may imply a Cmax component to this compound, although this will need to await more detailed PK-PD studies. The potential for a single IM injection to control *N. gonorrhoeae* is appealing, especially in clinical settings where repeated drug dosing and compliance may not be easy to achieve.

Finally, given the potent inhibitory activity of JSF-2414 against a range of high-threat Gram-positive organisms, a preliminary study was initiated to examine to explore the *in vivo* potential of JSF-2659 against MRSA and *C. difficile*. A murine deep-tissue thigh model was used to demonstrate a dose-dependent reduction of VRSA and maximum >6.2 log10 reduction to the near LOD when administered PO at 250 mg/kg TID (**Fig. 4**). Linezolid at 50 mg/kg PO BID and vancomycin at 100 mg/kg SC TID resulted in reductions of 3.41 and 2.09 log10, respectively. In a hamster colitis model, JSF-2659 administered PO at 20, 100 or 250 mg/kg BID for 5 days was fully efficacious in preventing *C. difficile* induced mortality over 14 days (max) relative to the untreated control which showed 100% mortality at day 4 (**Fig. 5**). Vancomycin at 20 mg/kg PO BID for 5 days was equally effective at preventing mortality.

In summary, the small molecule 8-(6-fluoro-8-(methylamino)-2-((2-methylpyrimidin-5- yl)oxy)-9H-pyrimido[4,5-b]indol-4-yl)-2-oxa-8-azaspiro[4.5]decan-3-yl)methanol developed as JSF-2414 is a tricyclic pyrimidoindole that was designed as a potent inhibitor of the ATP binding/hydrolysis region of DNA gyrase (GyrB) and topoisomerase (ParE). JSF- 2414 displays highly potent activity against the fastidious Gram-negative organism *N. gonorrhoeae* including fluoroquinolone-resistant and other drug-resistant strains To improve its oral bioavailability, a phosphate prodrug JSF-2659 was developed. Oral administration of JSF-2659 was highly efficacious against *N. gonorrhoeae* in reducing microbial burdens to the limit of detection.in two different animal models of vaginal infection, Furthermore, IM administration increased its PK T1/2 6-fold resulting in reduction in microbial burden to the LOD following a single dose. Lastly, due to its potent activity against Gram-positive organisms in vitro, JSF-2659 was shown in a preliminary preclinical deep tissue model of MRSA and model of *C. difficile*-induced colitis to be highly efficacious and protective. This new preclinical development candidate has strong potential to be used against high-threat multidrug resistant organisms. Its mechanism of action as a dual ATP-binder ensures that the development resistance will be extremely low.

## ACKNOWLEDGMENTS

Thomas Hiltke, Ph.D. Basic Research Program Officer Sexually Transmitted Infections Branch DMID/NIAID/NIH/DHHS and Kimberly Murphy, MS, Product Development Program Manager, STDB NIAID. This work is dedicated to the memory of Professor Roger Spanswick whose inspiration and courage helped move this program forward.

## DISCLAIMER

The opinions and assertions expressed herein are those of the author(s) and do not necessarily reflect the official policy or position of the Uniformed Services University or the Department of Defense. This work was prepared by a civilian employee (AEJ) of the US Government as part of the individual’s official duties and therefore is in the public domain and does not possess copyright protection.

## METHODS

### Bacterial strains

*N. gonorrhoeae* clinical isolates (n=100) were obtained as frozen stocks from the Centers for Disease Control and Prevention (CDC) ^33^. The strains were inventoried using the original designations and stored in a freezer set at -70°C; simple ID numbers (CDC 1-100) were assigned for internal use. The isolates were plated onto Chocolate II Agar (BD BBL™ # 221267) to assess purity and growth on solid medium at 36 ± 1°C in an atmosphere of 5% CO2. Plates were visually inspected for colony growth and morphology after 24 hours incubation. All isolates formed smooth, nonpigmented, small colonies characteristic for *N. gonorrhoeae. Staphylococcus epidermidis*, *Enterococcus faecium*, vancomycin-resistant enterococci (VRE), *E. faecalis*, *B. anthracis*, and a toxigenic *C. difficile* panel that included the common ribotypes 001, 002, 012, 014, 020, 038, 078, 087 and the highly virulent ribotype 027. Gram-negative pathogens included *Escherichia coli, Pseudomonas aeruginosa, Klebsiella pneumoniae* and *Acinetobacter baumannii* and select agents *F. tularensis and Y. pestis*.

### Susceptibility testing

#### Neisseria gonorrhoeae

Susceptibility testing was performed by the Southern Research Institute (Birmingham, AL) under NIH testing contract No: HHSN272201100012I. Six control antibacterials, azithromycin (Azi), cefixime (Cfx), ceftriaxone (Cro), ciprofloxacin (Cip), penicillin (Pen), and tetracycline (Tet), were purchased from commercial sources. Hundred-fold concentrated stock solutions of test compounds and control antibacterials were prepared in the appropriate solvent. DMSO was used for the test compound and for Azi, Cfx, Pen, Tet. The Cro stock solution was made with sterile water, and the Cip stock was made with 0.1 N HCl(aq). Potency values provided by the manufacturers of the drugs were used to calculate the amounts of powder needed to prepare stock solutions with a final concentration of 3.2 mg•ml^-1^ (Cip), 1.6 mg•ml^-1^ (Azi, Pen, Tet), and 0.4 mg•ml^-1^ (Cfx, Cro). The potency of the test compound powders was assumed to be 100% for the purpose of this study. The stock solutions of 50 µg•ml^-1^ compound were prepared in DMSO. The stock solutions were used as starting material for 2-fold serial dilutions in corresponding solvents to obtain compound solutions that could be directly added at a ratio of 1:100 (v/v %) to molten Gonococci agar (GC agar) before pouring rectangular assay plates with the same dimensions as the standard 96-well microplates (Thermo Scientific Nunc # 267060). The GC agar recommended for *N. gonorrhoeae* testing contains 3.6% GC agar base (Oxoid™ GC agar # CM0367) and 1% defined growth supplement (BD BBL™ IsoVitaleX # 211876). Compound solutions described above were incorporated into GC agar medium to pour rectangular plates that represent 2-fold serial dilutions consisting of 12 compound dilution steps of test or control article. The serial dilutions in agar medium were prepared in triplicate. JSF-2414 was tested in the range between 0.500 and .00024 µg•ml^-1^. For Azi, Pen, and Tet, the test range was from 16.00 to 0.008 µg•ml^-1^; Cfx and Cro were tested in the range between 4.00 and 0.002 µg•ml^-1^; Cip was assayed in the range between 32.00 and 0.016 µg•ml^-1^. Strains were grown on drug-free agar plates (BD BBL™ Chocolate II Agar plates) for 24 hours at 36 ± 1°C in 5% CO2. Colonies were suspended directly in 0.9% saline supplemented with 0.5x Mueller Hinton medium. In preparation for the MIC assay, the inocula were standardized by adjusting their turbidity to the 0.5 McFarland standard and transferred by dispensing 700-µl aliquots to 96-well deep well plates. The deep well plates with the inocula were kept on ice at all times. Replicators with 96 one-millimeter pins (Boekel Industries # 140500) were sterilized in an autoclave before the start of the experiment and with an open flame between the transfer steps. The use of multiple replicators ensured that the transfer pins had cooled down to ambient temperature before every transfer performed during the experiment. The 96-pin replicators were used to transfer 1 µl aliquots of bacterial suspension from the inocula-containing 96-well deep well plates to the agar surface of the assay plates containing test article or control antibiotic. The same method was used to inoculate compound-free agar plates for growth control. Plates were incubated for up to 48 hours at 36 ± 1°C in 5% CO2. After the incubation period, assay and growth control plates were inspected visually, and the MIC was recorded as the lowest concentration of compound that completely inhibited growth.

#### Bacterial culture

All of the strains of ESKAPE bacteria and *Burkholderia pseudomallei* was purchased from the American Type Cell Culture Institute (ATCC), Manassas, VA, USA. *Brucella melitensis* and *Yersinia pestis* were purchased from BEI Resources.

Mycobacterial strains were grown overnight in 7H9 broth (Becton, Dickinson and Company 271310), plus 0.2% glycerol (Sigma G5516), 0.25% Tween 80-20% (Sigma P8074) and 20% 5x ADC. The 5x ADC solution was prepared using 25 g/L Bovine Serum Albumin (Sigma A9647), 10 g/L dextrose (Sigma D9434) and 4.2 g/L NaCl (Sigma S5886). For the Minimum Inhibitory Concentration assay (MIC), all mycobacteria were grown in 7H9 without Tween. The strains of *Bacillus anthracis* and *Francisella tularensis* were obtained from the USAMRC Bacteriology Division, Ft. Detrick, MD, USA. *Enterococcus faecium* and *Staphylococcus epidermidis* were grown in BBL^TM^ Muller Hilton II cation adjusted broth (Becton, Dickinson and Company 212322) with the addition of 1% IsoVitaleX^TM^ (Becton, Dickinson and Company 211875). *Brucella melitensis* was grown in BBL^TM^ Brucella Broth (Becton, Dickinson and Company D 211088). *Francisella tularensis* was grown in Cysteine Heart Broth [10 g BBL^TM^ Brain Heart Infusion (Becton, Dickinson and Company 211059), 10 g Proteose Peptone (Sigma F29185), 10 g Dextrose (Sigma D9434), 5 g sodium chloride (Sigma S3014), 1g L-Cysteine (Sigma C7352) in 1 L water]. All of the other bacterial strains were grown in BBL^TM^ Muller Hilton II broth cation adjusted (MH).

#### Microsomal stability assay

Liver microsome stability assays were performed by BioDuro Inc. Human and mouse (CD-1 male) liver microsomes were used. Briefly, 2.5 µL of control compounds and test compound (dissolved and diluted in DMSO to 100 µM concentration) were added to 197.5 µL of reaction buffer (0.05 M Phosphate reaction buffer, pH=7.4) and vortexed. 50 µl of reaction buffer was added to solution and mixed via pipetting up and down 6 times. At each time point of 0, 30 and 60 min, an aliquot of 20 µL was removed from each tube. Liver microsomes (LM) working solution was prepared in 0.05 M Phosphate reaction buffer (pH=7.4) for final concentration of 1.27 mg/ml. NADPH solution was prepared in 0.05 M Phosphate buffer to afford a 5 mM buffer solution. 2.5 µL of positive control (5x mixed) and test compounds were added into 197.5 of LM working solution. After vortexing and incubating for 5 min (37°C), 50 µL of NADPH solution was added and mixed via pipetting. At each time point of 0, 5, 15, 30 and 60 min, an aliquot of 20 µL was removed from each tube. 250 µL of quenching solution was aliquoted to quench the reaction, which was then vortexed for 1 min, and placed on ice for 40 min. After centrifugation at 4,000 rpm for 15 min, 100 µL of supernatant was transferred using a multichannel pipette to 0.65 mL tubes. Samples were diluted with MeOH: H2O (1:1) as necessary. Samples were analyzed by high-pressure liquid chromatography coupled to tandem mass spectrometry (LC/MS/MS) performed on a Sciex Applied Biosystems Qtrap 4000 triple-quadrupole mass spectrometer coupled to a Shimadzu HPLC system to quantify the compound remaining. Data was processed using Analyst software (version 1.4.2; Applied Biosystems Sciex). Calculations were performed as CLint (µL/min/mg protein) = ln 2*1000 /T1/2 / Protein Conc., where the units of T1/2 were min, and the units of Protein Conc. were mg/mL. Metabolite identification on the microsomal extracts was performed using a Q-Exactive HRMS coupled with an Ultimate 3000 HPLC system and a Kinetix C18 2.1x50mm 2.6um HPLC column.

### Animal Models

#### N. gonorrhoeae vaginal infection models

Efficacy of JSF-2659 in clearing an experimental *N. gonorrhoeae* infection of MDR strain H041 in female mice was performed by Dr. Ann Jerse’s laboratory at the Uniformed Services University (USU). Four dosing regimens of JSF-2659 were tested as described in **Supplement, Fig. 1.** A total of 60 female BALB/c mice (6 groups; n = 8-10 mice/group) were implanted with a 21-day slow-release, 5 mg 17-β estradiol pellet (Innovative Research of America) under the skin (day -4) to induce susceptibility to *N. gonorrhoeae*. Antibiotics (streptomycin [STM], trimethoprim sulfate [TMP]), were administered to suppress the overgrowth of commensal flora that occurs under the influence of estradiol ^17, 34^. Mice were inoculated on day -2 (two days after estradiol treatment was initiated) with a dose of the challenge strain that infects 80 – 100% of mice (ID80-100) (10^4^ CFU for strain H041(STM^R^), a streptomycin-resistant derivative of the ceftriaxone resistant strain H041, referred to as H041 for brevity). Strain H041 was selected because of its clinical significance as a multidrug resistant ‘superbug’ ^18^. Vaginal mucus was quantitatively cultured for *N. gonorrhoeae* on the morning of the next two consecutive days (days -1 and 0) to confirm infection. On the day of treatment (day 0), JSF-2659 was solubilized in 0.5% Carboxymethyl Cellulose Sodium Salt (CMC; MP Bio) and 0.5%Tween 80 (Fisher) in sterile endotoxin-free water; this vehicle solution was also used as the negative control.

On the afternoon of day 0 (and following the morning culture), JSF-2659 was administered orally (PO) as either a single dose (QD) or three doses every 6 hours (TID) over a 24 h period. The vehicle control was administered TID. The positive control that was established for H041 was 5 doses of 48 mg/kg gentamicin (GEN) administered once daily for 5 days IP (0.2 mL), with treatment beginning on day 0 ^35^. Experimental groups were designated as outlined in **Supplement, Fig. 1.** Vaginal mucus was quantitatively cultured for *N. gonorrhoeae* for 8 consecutive days after treatment (days +1 through +8). Vaginal material was collected by wetting a Puritan rayon swab in sterile, endotoxin-free PBS, gently inserting the swab into the vagina, and suspending the swab in 1 mL of GCB. Broth suspensions were diluted in GCB (1:100 for H041), and diluted and undiluted samples were cultured on GC-VCNTS agar using the Autoplater automated plating system (Spiral Biotech). GC-VCNTS agar contained vancomycin, colistin, nystatin, trimethroprim sulfate (VCNT supplement; Difco BD, Product #202408) and 100 µg/mL streptomycin sulfate. A portion of the swab was also inoculated onto heart infusion agar (HIA) to monitor the presence of facultative aerobic commensal flora. The number of viable *N. gonorrhoeae* bacteria recovered were determined using the Spiral Biotech Q- Counter Software at 48 h of incubation at 37 °C. The percentage of mice in each test group that were culture positive at each time point were plotted as Kaplan Meier colonization curves and compared to the positive control and vehicle control, and the other test groups. Differences were analyzed by the Log-rank (Mantel-Cox) test. Colonization load, defined as the number of CFU per ml of vaginal swab suspension, were also compared among groups using a repeated measures ANOVA with Bonferroni multiple comparisons. *p* values < 0.05 were considered significant. Mice must have had at least 3 consecutive days of negative culture to be considered cleared of infection. At the study endpoint, mice were euthanized using compressed CO2 gas in a CO2 gas chamber in the Laboratory Animal Medicine Facility. All animal experiments were conducted at the Uniformed Services University of the Health Sciences, a facility fully accredited by the Association for the Assessment and Accreditation of Laboratory Animal Care, under a protocol that was approved by the USUHS Institutional Animal Care and Use Committee in accordance with all applicable Federal regulations governing the protection of animals in research.

A secondary model (Pharmacology Discovery Services Taiwan, Ltd) was performed according to Song et al. ^36^. Vaginal infection model using *N. gonorrhoeae* strain F1090 (ATCC 700825) was performed with groups of 5 ovariectomized BALB/c mice aged 5-6 weeks. Ovariectomy was performed at 4 weeks of age. The period of surgical recovery and acclimation was ∼7 days. Animals were subcutaneously (SC) injected with 17β- estradiol solution solubilized in cotton seed oil at 0.23 mg/mouse 2 days before infection (Day -2) and on the day of infection (Day 0). To minimize the commensal vaginal bacteria, animals were treated twice daily (BID) with streptomycin (1.2 mg/mouse) and vancomycin (0.6 mg/mouse) by IP injection along with trimethoprim sulfate at 0.4 mg/mL supplied in the drinking water. Antibacterial treatments were started two days prior to infection and were continued daily until the end of study. On Day 0, animals were inoculated intravaginally (IVG) with *N. gonorrhoeae* under anesthesia with IP injection of pentobarbital (80 mg/kg). The vagina was rinsed with 50 mM Hepes (pH 7.4, 30 μL) followed by inoculation with *N. gonorrhoeae* ATCC 700825 suspension, 0.02 mL/mouse. The target inoculation density was 2.0 × 10^6^ CFU/mouse and the actual count was 1.14 x 10^6^ CFU/mouse. A 0.05 mL aliquot was inoculated into a chocolate agar plate and incubated at 35-37°C with 5% CO2 overnight. The culture was re-suspended in 1 mL PBS (>1.0 x 10^10^ CFU/mL, OD620 2.0 – 2.2) and diluted in PBS containing 0.5 mM CaCl2 and 1 mM MgCl2 to obtain the target inoculum of 1.0 x 10^8^ CFU/mL. Colony counts were determined by plating dilutions to chocolate agar plates followed by 20 – 24 h incubation. The actual CFU count was 5.7 x 10^7^ CFU/mL. JSF-2659 at 75 and 250 mg/kg, was administered orally (PO) with two dosing schedules, including once (QD) at 2 h after infection, and three times (TID) with a 6 h interval at 2, 8, and 14 h after infection. Test articles were also administered intramuscularly (IM) with JSF-2659 at 75 and 250 mg/kg once (QD) at 2 h after infection. Two reference compounds were applied in this study, ciprofloxacin and gentamicin. Ciprofloxacin was administered orally (PO) at 12.5 mg/kg once (QD) at 2 h after infection. Gentamicin was administered intraperitoneally (IP) at 48 mg/kg once (QD) at 2 h after infection for 1, 3 or 7 consecutive days depending on the time point of animal scarification, at 26, 74 or 170 h after infection. One infected but non-treated group was sacrificed at 2 h after infection for the initial bacterial counts. Each animal was weighed prior to each dose and the dose volumes were 10 mL/kg for all dosing groups. Test animals were euthanized by CO2 asphyxiation and sacrificed at 26, 74 or 170 h after infection. Vaginal lavage was performed twice with 200 μL GC broth containing 0.05% saponin to recover vaginal bacteria and the lavage fluids were pooled in a total volume of 500 μL. The bacterial counts (CFU/mL) in lavage fluid were calculated and the percentage decrease relative to the vehicle control was calculated by the following formula: Decrease (%) = [(CFU/mL of vehicle – CFU/mL of treatment) / (CFU/mL of vehicle)] x 100%. Statistical significance was assessed with one-way ANOVA followed by Dunnett’s method using the Prism Graphpad software version 5.0. A significant (*p* < 0.05) decrease in the bacterial counts of the treated animals compared to the vehicle control group was considered significant difference.

### Staphylococcus soft tissue infection model

A deep tissue thigh model of *Staphylococcus aureus* was contracted to Pharmacology Discovery Services Taiwan, Ltd. All studies were performed with *S. aureus* strain VRS-2 – a VRSA strain Hershey, Van-A producing SCC Mec II, st5 strain that was isolated from the foot ulcer of a 70-year-old patient. It is methicillin-resistant with resistance to carbapenems, cephalosporins and penicillins. VRS-2 is resistant to vancomycin (MIC >64 µg/ml), quinolones (LVX-R, CIP- R), macrolides (ERY-R, CLI-R), and trimethoprim/sulfamethoxazole. Groups of 5 male or female specific-pathogen-free ICR mice weighing 22 ± 2 g were used. Animals were immunosuppressed by two intraperitoneal injections of cyclophosphamide, the first at 150 mg/kg 4 days before infection (Day –4) and the second at 100 mg/kg 1 day before infection (Day –1). On day 0, animals were inoculated intramuscularly (0.1 ml/thigh) with *Staphylococcus aureus*, vancomycin resistant (VRS-2) into the right thigh. Vehicle and/or test substances including linezolid at 50 mg/kg BID were then administered (SC or PO) 2 and 14 hours later. At 24 hours after treatment, animals were humanely euthanized with CO2 asphyxiation and then the muscle of the right thigh was harvested from each test animal. The removed muscle tissues were homogenized in 3 ml of PBS, pH 7.4, with a polytron homogenizer. Homogenates, 0.1 ml, were used for serial 10-fold dilutions and plated on nutrient agar plates for colony count

### Hamster Colitis Model

This model contracted to Pharmacology Discovery Services Taiwan, Ltd assessed test articles for protection against a lethal *C. difficile* colitis infection ^37^. Groups of 10 male or female Golden Syrian hamsters weighing 90 ± 10 g were used.

Each animal was pretreated with a single subcutaneous dose of clindamycin at 50 mg/kg (Day -1) to induce susceptibility to *C. difficile*. Twenty-four hours after the clindamycin treatment (Day 0), the animals were infected with *C. difficile* BAA-1805, a ribotype 027 strain. Spores were administered in a single oral lethal (LD90-100) dose, 1 × 10^5^ spores per animal. Test substance and vehicle were administered (PO, IP, IV, or SC) 16 hours after inoculation then twice daily (bid) for a total of 5 consecutive days. The mortality was recorded daily for 14 days following infection. Prevention of mortality in 50 percent or more of the animals indicated significant activity. *C. difficile* strain BAA-1805, toxigenic, NAP1, Ribotype 027 strain, was obtained from the American Type Culture Collection (Rockville, MD, USA) and cryopreserved as single-use frozen working stock cultures stored at -80°C. A 0.1 mL aliquot was transferred to anaerobic blood agar plates (anaerobic BAP) and incubated in an anaerobic workstation (Don Whitley A35) at 35- 37°C anaerobic condition (80% N2, 10% CO2, 10% H2) for 5 days. Growth on plates was transferred to phosphate buffer saline (PBS) and heated at 70°C for 30 minutes to inactivate the spores. The heated culture was pelleted by centrifugation at 3,500 x g for 15 minutes, and then re-suspended in cold PBS (>5 x 10^7^ spores/mL in original). The culture was diluted in PBS to an estimated concentration of 0.6-1.0 x 10^6^ spores/mL. The actual colony counts were determined by plating dilutions onto CCFA-HT plates followed by 2 days incubation and colony counting. The actual inoculum was 1.16 x 10^6^ spores/mL. Male golden Syrian hamsters 6 weeks of age were provided by National Laboratory Animal Center, Taiwan. The animals were individually housed in animal cages. All animals were maintained in a well-controlled temperature (20 - 24°C) and humidity (30% - 70%) environment with 12 hours light/dark cycles. Free access to standard lab diet [MFG (Oriental Yeast Co., Ltd., Japan)] and autoclaved tap water were granted for study period. All aspects of this work including housing, experimentation, and animal disposal were performed in general accordance with the Guide for the Care and Use of Laboratory Animals (National Academies Press, Washington, D.C., 2011). The study was performed in our AAALAC accredited ABSL2 laboratory under the supervision of staff veterinarians. The animal care and use protocol was approved by the IACUC of Pharmacology Discovery Services Taiwan Ltd.

### Quantitative Methods

Levels of JSF-2414 in plasma and tissues were measured by LC-MS/MS in electrospray positive-ionization mode (ESI+) on a Sciex Qtrap 4000 triple-quadrupole mass combined with an Agilent 1260 HPLC using Analyst software and multiple-reaction monitoring (MRM) of precursor/product transitions. The following MRM transitions were used for JSF-2414 (494.28/324.80) and the internal standard Verapamil (455.4/165.2). Chromatography was performed with an Agilent Zorbax SB-C8 column (2.1x30 mm; particle size, 3.5µm) using a reverse phase gradient elution. 0.1% formic acid in Milli-Q deionized water was used for the aqueous mobile phase and 0.1% formic acid in acetonitrile (ACN) for the organic mobile phase. Tissues were homogenized prior to extraction by combining 4 parts PBS buffer: 1 part tissue. The samples were homogenized using a SPEX Sample Prep Geno/Grinder 2010 for 5 minutes at 1500 RPM. 1 mg/mL DMSO stock was serial diluted in 50/50 ACN/water to create standard curves and quality control spiking solutions. 20 µL of neat spiking solutions were added to 20 µL of drug free mouse K2EDTA plasma (Bioreclammation) or drug free tissue homogenate and extraction was performed by adding 200 µL of acetonitrile/methanol 50/50 protein precipitation solvent containing 10 ng/mL Verapamil (Sigma). Extracts were vortexed for 5 minutes and centrifuged at 4000 RPM for 5 minutes. The supernatants were analyzed by LC-MS. Sample analysis was accepted if the concentrations of the quality control samples were within 20% of the nominal concentration.

## SUPPLEMENT

**Figure S1.**
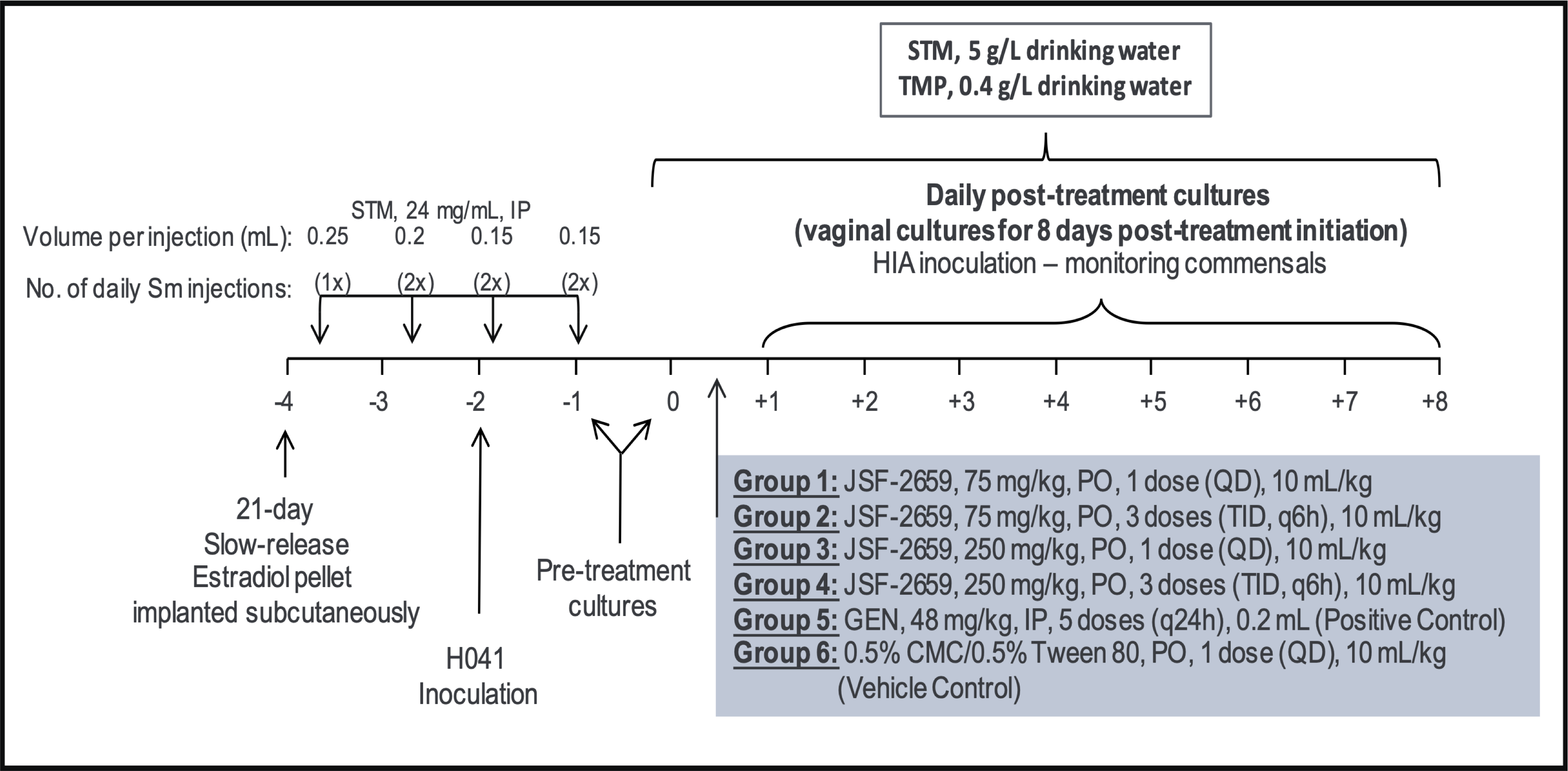
Experimental design schematic of the *Neisseria gonorrhoeae* infection model with strain H041. The following test regiments were performed: JSF-2659; <u>Group 1</u>: 75 mg/kg, PO, 1 dose (QD), 10 mL/kg; <u>Group 2</u>: 75 mg/kg, PO, 3 doses every 6 h (TID), 10 mL/kg; <u>Group 3</u>: 250 mg/kg, PO, 1 dose (QD), 10 mL/kg; <u>Group 4</u>: 250 mg/kg, PO, 3 doses every 6 h (TID),10 mL/kg; Positive control group <u>Group 5:</u> GEN, 48 mg/kg, 5 doses once daily, IP, 0.2 mL and Vehicle control group <u>Group 6</u>: 0.5% CMC/0.5% Tween 80, 3 doses (TID), 10 mL/kg

## Methods

### Preparation of JSF-2414 and JSF-2659

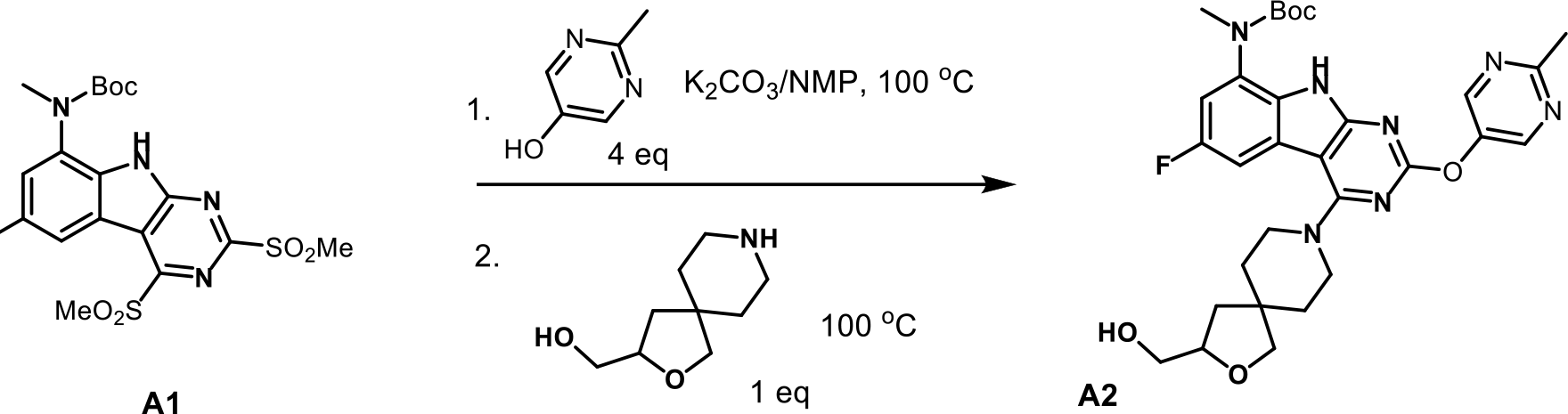

To *tert*-butyl (6-fluoro-2,4-bis(methylsulfonyl)-9H-pyrimido[4,5-b]indol-8- yl)(methyl)carbamate (9.45 g, 20.0 mmol; Source: WuXiAppTec) in NMP (50 mL) was added 2-methylpyrimidine-5-ol (8.80 g, 80.0 mmol) and potassium carbonate (11.00 g, 79.7 mmol). The mixture was heated at 100 °C for 70 min, then (-)-(2-oxa-8- azaspiro[4.5]decan-3-yl)methanol hydrogen chloride salt(4.15 g, 20.0 mmol; Source: WuXiAppTec; [α]D22 = -3.14 (c 0.1, MeOH)) was added. The mixture was heated at 100°C for 2.5 h. 400 mL water was added with stirring, after which the precipitate was filtered and washed with water to give *tert*-butyl (6-fluoro-4-(3-(hydroxymethyl)-2-oxa-8- azaspiro[4.5]decan-8-yl)-2-((2-methylpyrimidin-5-yl)oxy)-9H-pyrimido[4,5-b]indol-8- yl)(methyl)carbamate as a yellow solid in 71.5% yield (8.50 g, 14.3 mol). The product was used in the next step without further purification.

### Scheme 2

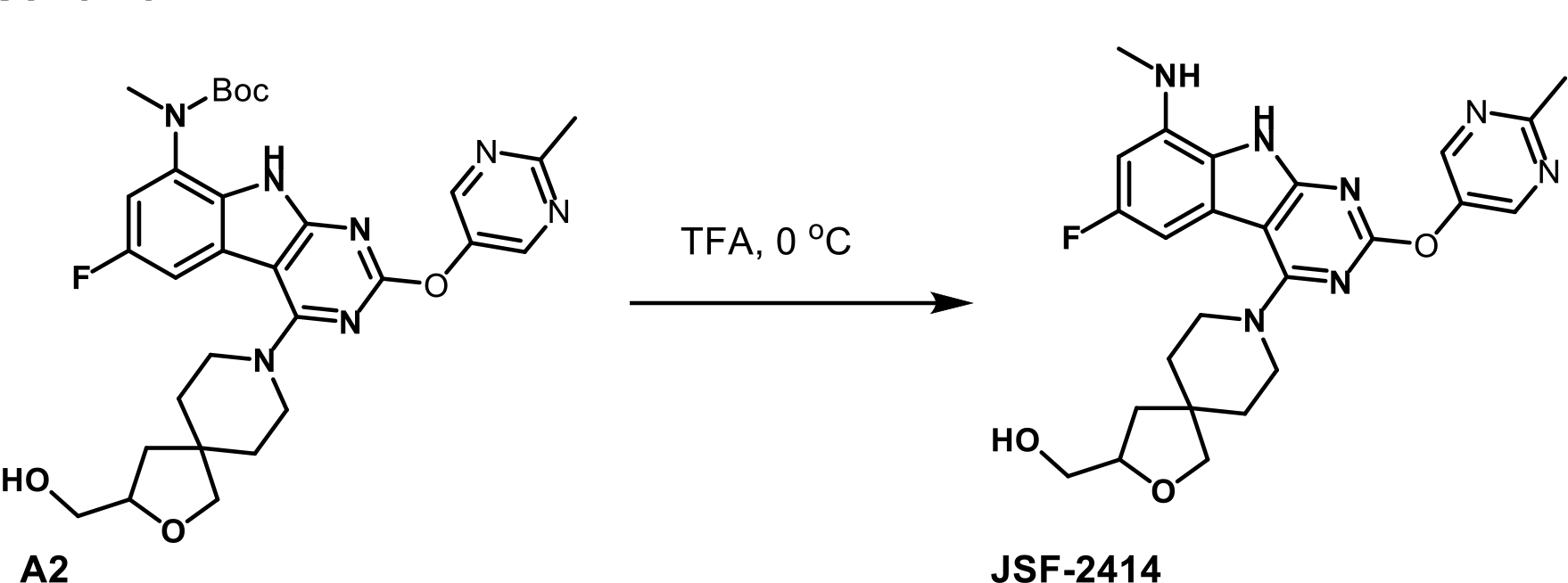

A mixture of A2 = *tert*-butyl (6-fluoro-4-(3-(hydroxymethyl)-2-oxa-8- azaspiro[4.5]decan-8-yl)-2-((2-methylpyrimidin-5-yl)oxy)-9H-pyrimido[4,5-b]indol-8- yl)(methyl)carbamate (3.00 g, 5.05 mmol) in 12 mL TFA was stirred at 0 °C for 1 min. The mixture was concentrated *in vacuo*, after which it was diluted with 10 mL ethanol and 20 mL water. The pH of the mixture was adjusted to 10 by adding 6 N NaOH(aq).The resulting mixture was stirred for 8 h and was then poured into 50 mL saturated NH4Cl(aq). The mixture was extracted with ethyl acetate (4x50 mL). The combined extracts were diluted to 100 mL and washed with saturated aqueous brine solution. The organic layer was dried and concentrated under reduced pressure. Purification by flash column chromatography on silica gel, eluting with 0-5% methanol/dichloromethane provided JSF- 2414 as a yellow solid in 73.9% yield (1.84 g, 3.73 mol). ^1^H NMR (500 MHz, d6-DMSO) δ 11.7 (s, 1), 8.73 (s, 2), 6.63 (dd, J = 10.0, 1.3 Hz, 1), 6.35 (dd, J = 12.1, 1.4 Hz, 1), 5.58 (d, J = 2.9 Hz, 1), 4.67 (s, 1), 4.04 – 3.90 (m, 1), 3.68 (ddd, J = 18.1, 13.0, 5.1 Hz, 2), 3.63 – 3.47 (m, 4), 3.41 (d, J = 4.7 Hz, 2), 2.85 (d, J = 3.8 Hz, 3), 2.66 (s, 3), 1.93 (dd, J = 12.3, 7.2 Hz, 1). [α]D22 = -3.49 (c 0.5, MeOH). LRMS m/z: [M+H]^+^Calcd for C25H29FN7O3 494.2; found 494.2.

#### Scheme 3

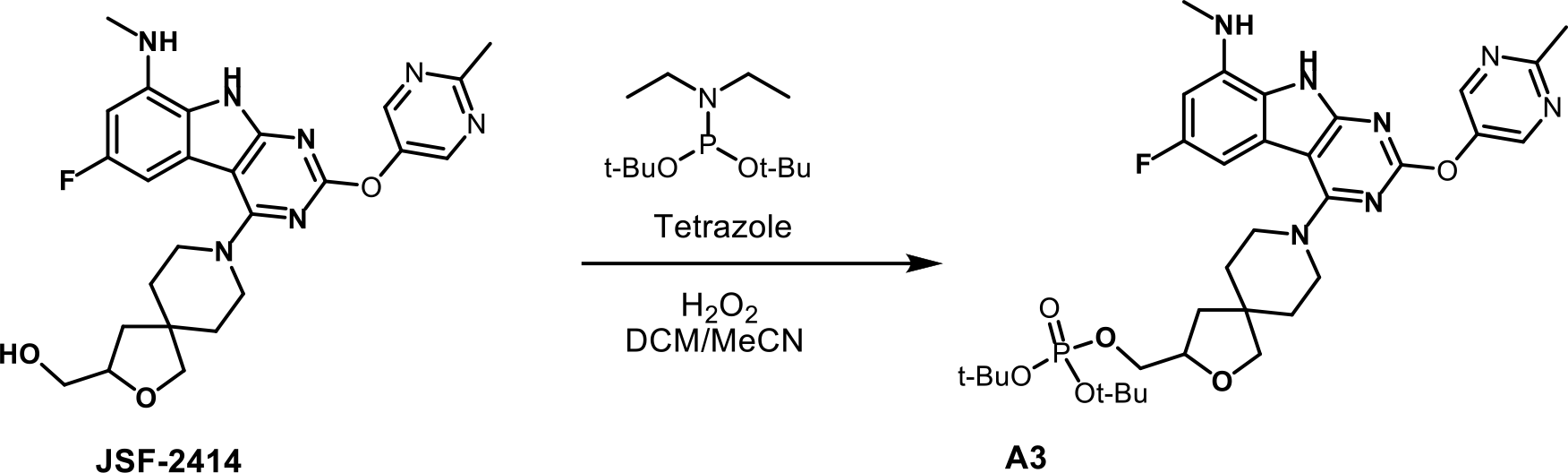

To JSF-2414 (1.45 g, 2.94 mmol) in dichloromethane (20 mL) and acetonitrile (2 mL) was added a tetrazole solution (18.0 mL, 8.10 mmol, 0.45 M in acetonitrile) and di- *tert*-butyl N,N-diethylphosphoramidite (2.20 mL, 7.91 mmol). The mixture was stirred at rt for 2 h. Then a 20% aqueous solution of H2O2 (3.0 mL) was added. After being stirred at rt for 10 min, the mixture was extracted with ethyl acetate. The organic phase was washed with saturated aqueous brine solution and dried over anhydrous sodium sulfate. The solution was filtered, concentrated and purified by flash column chromatography on silica gel, eluting with 0 – 4% methanol/dichloromethane to give di-*tert*-butyl ((8-(6-fluoro-8- (methylamino)-2-((2-methylpyrimidin-5-yl)oxy)-9H-pyrimido[4,5-b]indol-4-yl)-2-oxa-8- azaspiro[4.5]decan-3-yl)methyl) phosphate as a brown oil in 64.6% yield (1.30 g, 1.90 mmol). LRMS m/z: [M+H]^+^Calcd for C33H6FN7O3P 686.3; found 686.2.

#### Scheme 4

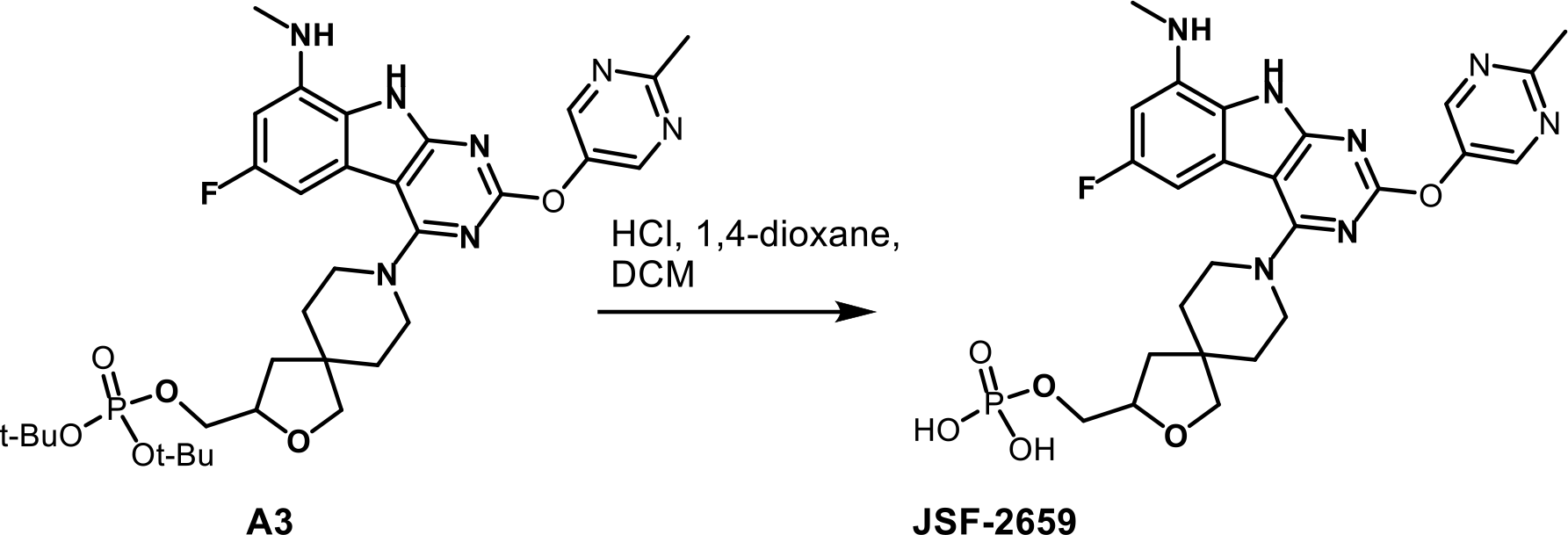

To A3 = di-*tert*-butyl ((8-(6-fluoro-8-(methylamino)-2-((2-methylpyrimidin-5-yl)oxy)- 9H-pyrimido[4,5-b]indol-4-yl)-2-oxa-8-azaspiro[4.5]decan-3-yl)methyl) phosphate (260 mg, 0.379 mmol) in dichloromethane (10 mL) was added 0.7 mL 4 N HCl(aq) in 1,4-dioxane (2.8 mmol) dropwise. The mixture was stirred at rt for 10 min. Then dichloromethane was removed by pipette. The solid was washed with ethyl acetate (2 x 10 mL) and dried *in vacuo* to give JSF-2659 as a yellow powder in 99% yield (250 mg, 0.377 mmol). Elemental analysis was consistent with two HCl and one H2O molecules per molecule of targeted product. ^1^H NMR (500 MHz, D2O) δ 8.53 (s, 2), 6.27 (d, J = 11.8 Hz, 1), 6.22 (d, J = 10.2 Hz, 1), 4.26 – 4.14 (m, 1), 3.82 – 3.72 (m, 1), 3.70 – 3.61 (m, 1), 3.58 (s, 2), 3.29 – 3.05 (m, 4), 2.75 (s, 3), 2.62 (s, 3), 1.97 – 1.85 (m, 1), 1.46 (s, 5). Two Hs were unaccounted for and presumably were the two N-Hs which were exchanging with D2O. LRMS m/z: [M+H]^+^Calcd for C25H29FN7O6P 573.2; found 574.2. The bis-sodium salt JSF-2659-B was prepared by adding saturated aqueous NaHCO3 to the HCl salt in water. Purification by HPLC, eluting with water/acetonitrile gave a white solidJSF-2659-B. 1H NMR (500 MHz, d6-DMSO) δ 11.9 (s, 1), 8.76 (s, 2), 6.67 (d, J = 9.8 Hz, 1), 6.41 (d, J = 11.8, 1), 4.10 – 4.10 (m, 1), 3.87 – 3.77 (m, 2), 3.75 – 3.62 (m, 2), 3.59 (s, 3), 3.57 – 3.49 (m, 2), 2.85 (s, 3), 2.67 (s, 3), 1.73 – 1.61 (m, 4), 1.59(s, 1), 1.53 (dd, J = 12.2, 8.6 Hz, 1). Also noted 10.3– 9.0 (brs, solvent H2O bound to OH of acid). LRMS m/z: [M+H]^+^Calcd for C25H30FN7O6P 574.2; found 574.2.

**Table 1.** Individual MIC values for clinical isolates of *Neisseria gonorrhoeae*

| Strain ID | | Test and Control Articles MIC ($\mu\text{g}\cdot\text{ml}^{-1}$) | | | | | | |
| --- | --- | --- | --- | --- | --- | --- | --- | --- |
| Southern Research | CDC | JSF-2414 | Azi | Cfx | Cro | Cip | Pen | Tet |
|  |  | MIC Range | MIC Range | MIC Range | MIC Range | MIC Range | MIC Range | MIC Range |
|  |  | 0.0005-0.03 | 0.016->16.0 | 0.016 – 4 | <0.002-0.125 | <0.016 – 16 | <0.008 – 4 | 0.063 – 16 |
| CDC1 | GCREF2012016 | 0.0039 | 0.25 | 0.125 | 0.031 | 8 | 1 | 2 |
| CDC2 | GCREF2012045 | 0.0039 | 0.5 | 0.125 | 0.031 | 16 | 1 | 1 |
| CDC3 | GCREF2013013 | 0.0039 | 0.125 | 0.125 | 0.031 | 8 | 1 | 1 |
| CDC4 | GCREF2013030 | 0.0039 | 0.125 | 0.125 | 0.031 | 8 | 1 | 0.5 |
| CDC5 | GCREF2012001 | 0.0039 | 0.5 | 0.125 | 0.063 | 8 | 1 | 1 |
| CDC6 | GCREF2012002 | 0.0078 | 0.5 | 0.25 | 0.063 | 8 | 1 | 1 |
| CDC8 | GCREF2012004 | 0.002 | 0.125 | 0.125 | 0.016 | 8 | 0.25 | 1 |
| CDC9 | GCREF2012005 | 0.0039 | 0.125 | 0.125 | 0.031 | 8 | 1 | 1 |
| CDC10 | GCREF2012006 | 0.0039 | 0.5 | 0.125 | 0.031 | 16 | 1 | 1 |
| CDC11 | GCREF2012007 | 0.0039 | 0.25 | 0.125 | 0.031 | 8 | 1 | 1 |
| CDC12 | GCREF2012008 | 0.0039 | 0.125 | 0.25 | 0.031 | 8 | 1 | 1 |
| CDC13 | GCREF2012009 | 0.0039 | 0.125 | 0.063 | 0.031 | 8 | 1 | 1 |
| CDC14 | GCREF2012010 | 0.0039 | 0.125 | 0.25 | 0.031 | 8 | 1 | 1 |
| CDC15 | GCREF2012011 | 0.0078 | 4 | 0.031 | 0.008 | 2 | 0.125 | 0.5 |
| CDC16 | GCREF2012012 | 0.0078 | 0.25 | 4 | 0.063 | 8 | 2 | 1 |
| CDC18 | GCREF2012014 | 0.0078 | 0.5 | 0.063 | 0.063 | 16 | 1 | 1 |
| CDC19 | GCREF2012015 | 0.0039 | 8 | 0.125 | 0.008 | <0.016 | 0.25 | 0.5 |
| CDC20 | GCREF2012017 | 0.016 | >16.0 | 0.125 | 0.063 | 2 | 0.125 | 1 |
| CDC21 | GCREF2012018 | 0.0039 | 0.125 | 0.25 | 0.031 | 16 | 1 | 1 |
| CDC22 | GCREF2012019 | 0.03 | 0.5 | 0.031 | 0.008 | 4 | 0.25 | 1 |
| CDC23 | GCREF2012020 | 0.0078 | 0.125 | 0.125 | 0.031 | 8 | 1 | 1 |
| CDC24 | GCREF2012021 | 0.0039 | 0.25 | 0.063 | 0.063 | 8 | 1 | 1 |
| CDC25 | GCREF2012022 | 0.0039 | 0.125 | 0.25 | 0.031 | 8 | 2 | 1 |
| CDC26 | GCREF2012023 | 0.0078 | 0.5 | 0.063 | 0.031 | 4 | 0.25 | 0.5 |
| CDC27 | GCREF2012024 | 0.0039 | 0.25 | 0.063 | 0.031 | 8 | 1 | 1 |
| CDC28 | GCREF2012025 | 0.0039 | 0.25 | 0.25 | 0.031 | 16 | 1 | 1 |
| CDC29 | GCREF2012026 | 0.0078 | 0.5 | 0.125 | 0.016 | <0.016 | 1 | 1 |
| CDC30 | GCREF2012027 | 0.0078 | 0.25 | 0.125 | 0.031 | 8 | 1 | 1 |
| CDC31 | GCREF2012028 | 0.0078 | 0.5 | 0.25 | 0.031 | 8 | 2 | 2 |
| CDC32 | GCREF2012029 | 0.0078 | 0.125 | 0.125 | 0.063 | 8 | 1 | 0.5 |
| CDC33 | GCREF2012030 | 0.0078 | 0.125 | 0.5 | 0.125 | 2 | 1 | 0.5 |
| CDC34 | GCREF2012031 | 0.0039 | 0.125 | 0.063 | 0.125 | 16 | 1 | 1 |
| CDC35 | GCREF2012032 | 0.0039 | 0.25 | 0.25 | 0.125 | 16 | 2 | 2 |
| CDC36 | GCREF2012033 | 0.002 | 4 | 0.063 | 0.031 | 8 | 0.25 | 1 |
| CDC37 | GCREF2012034 | 0.0039 | 0.25 | 0.125 | 0.031 | 16 | 1 | 1 |
| CDC38 | GCREF2012035 | 0.0039 | 0.25 | 0.063 | 0.031 | 16 | 1 | 1 |
| CDC39 | GCREF2012036 | 0.0039 | 0.25 | 0.25 | 0.031 | 8 | 2 | 1 |
| CDC40 | GCREF2012037 | 0.0039 | 0.25 | 0.125 | 0.031 | 8 | 1 | 1 |
| CDC41 | GCREF2012038 | 0.0039 | 0.25 | 0.25 | 0.031 | 8 | 2 | 1 |
| CDC42 | GCREF2012039 | 0.0039 | 0.25 | 0.25 | 0.031 | 16 | 2 | 2 |
| CDC43 | GCREF2012040 | 0.0039 | 0.25 | 0.25 | 0.031 | 16 | 2 | 2 |
| CDC45 | GCREF2012042 | 0.0078 | 0.5 | 0.25 | 0.125 | 16 | 2 | 2 |
| CDC46 | GCREF2012043 | 0.0039 | 0.125 | 0.125 | 0.031 | 16 | 1 | 1 |
| CDC47 | GCREF2012044 | 0.0039 | 0.5 | 0.125 | 0.031 | 16 | 1 | 1 |
| CDC48 | GCREF2012046 | 0.0039 | 0.25 | 0.125 | 0.031 | 8 | 1 | 1 |
| CDC49 | GCREF2012047 | 0.0039 | 0.5 | 0.25 | 0.031 | 16 | 2 | 1 |
| CDC50 | GCREF2012048 | 0.0078 | 0.25 | 0.25 | 0.063 | 16 | 2 | 1 |
| CDC51 | GCREF2012049 | 0.0039 | 0.125 | 0.125 | 0.008 | 16 | 2 | 1 |
| CDC52 | GCREF2012050 | 0.0039 | 0.125 | 0.25 | 0.008 | 16 | 2 | 1 |
| CDC53 | GCREF2012051 | 0.0078 | 0.125 | 0.25 | 0.004 | 16 | 2 | 1 |
| CDC54 | GCREF2012052 | 0.0039 | 0.016 | 0.25 | 0.004 | 16 | 2 | 1 |
| CDC55 | GCREF2013001 | 0.0078 | 0.125 | 0.125 | 0.031 | 16 | 1 | 1 |
| CDC56 | GCREF2013002 | 0.0039 | 0.125 | 0.25 | 0.016 | 16 | 1 | 1 |
| CDC57 | GCREF2013003 | 0.0039 | 0.125 | 0.125 | 0.004 | <0.016 | 1 | 0.5 |
| CDC58 | GCREF2013004 | 0.002 | 0.125 | 0.25 | 0.004 | 16 | 2 | 1 |
| CDC59 | GCREF2013044 | 0.002 | 0.125 | 0.25 | 0.004 | 4 | 2 | 16 |
| CDC60 | GCREF2013005 | 0.0078 | 0.125 | 0.125 | 0.031 | 16 | 4 | 1 |
| CDC61 | GCREF2013006 | 0.0078 | 0.25 | 0.125 | 0.031 | 32 | 4 | 1 |
| CDC62 | GCREF2013007 | 0.016 | 0.125 | 0.25 | 0.008 | 0.125 | 0.25 | 0.5 |
| CDC63 | GCREF2013008 | 0.0078 | 0.016 | 0.125 | <0.002 | <0.016 | <0.008 | 0.063 |
| CDC64 | GCREF2013009 | 0.016 | 0.125 | 0.125 | 0.008 | <0.016 | 0.5 | 1 |
| CDC65 | GCREF2013010 | 0.0078 | 0.125 | 0.125 | 0.016 | <0.016 | 0.5 | 0.5 |
| CDC66 | GCREF2013011 | 0.016 | 4 | 0.125 | 0.008 | <0.016 | 0.25 | 0.5 |
| CDC67 | GCREF2013014 | 0.0078 | 0.125 | 0.125 | 0.008 | 0.25 | 0.5 | 1 |
| CDC68 | GCREF2013016 | 0.002 | 0.125 | 0.063 | 0.016 | 16 | 1 | 1 |
| CDC69 | GCREF2013017 | 0.002 | 0.125 | 0.063 | 0.031 | 16 | 1 | 1 |
| CDC70 | GCREF2013018 | 0.0039 | 4 | 0.125 | 0.004 | 16 | 2 | 1 |
| CDC71 | GCREF2013019 | 0.016 | 0.125 | 0.016 | 0.008 | 0.25 | 0.25 | 1 |
| CDC72 | GCREF2013020 | 0.0078 | 0.125 | 0.25 | 0.031 | 16 | 2 | 1 |
| CDC73 | GCREF2013021 | 0.016 | 0.125 | 0.125 | 0.008 | 0.125 | 0.5 | 1 |
| CDC74 | GCREF2013022 | 0.016 | 0.125 | 0.125 | 0.008 | <0.016 | 0.25 | 1 |
| CDC75 | GCREF2013023 | 0.0078 | 0.5 | 0.125 | 0.016 | <0.016 | 0.5 | 1 |
| CDC76 | GCREF2013024 | 0.0078 | 0.125 | 0.125 | 0.016 | <0.016 | 0.5 | 1 |
| CDC77 | GCREF2013025 | 0.0078 | 4 | 0.125 | 0.008 | 0.25 | 0.125 | 1 |
| CDC78 | GCREF2013026 | 0.0005 | 0.031 | 0.063 | 0.016 | 2 | 0.5 | 1 |
| CDC79 | GCREF2013027 | 0.0039 | 0.125 | 0.125 | 0.016 | 16 | 1 | 1 |
| CDC80 | GCREF2013028 | 0.0039 | 8 | 0.125 | 0.008 | 0.25 | 2 | 1 |
| CDC81 | GCREF2013029 | 0.0039 | 0.125 | 0.25 | 0.031 | 16 | 2 | 1 |
| CDC82 | GCREF2013031 | 0.0078 | 2 | 0.125 | 0.031 | 32 | 1 | 0.5 |
| CDC83 | GCREF2013032 | 0.016 | 0.125 | 0.125 | 0.008 | 0.125 | 0.25 | 0.5 |
| CDC84 | GCREF2013033 | 0.0078 | 0.125 | 0.125 | 0.008 | <0.016 | 0.25 | 0.5 |
| CDC86 | GCREF2013035 | 0.0078 | 0.125 | 0.125 | 0.008 | 0.25 | <0.008 | 0.25 |
| CDC87 | GCREF2013036 | 0.0039 | 0.125 | 0.25 | 0.063 | 16 | 2 | 1 |
| CDC88 | GCREF2013037 | 0.0078 | 2 | 0.125 | 0.031 | 16 | 2 | 0.5 |
| CDC89 | GCREF2013038 | 0.0039 | 2 | 0.125 | 0.008 | 16 | 1 | 1 |
| CDC90 | GCREF2013039 | 0.0078 | 0.125 | 0.125 | 0.031 | 16 | 2 | 1 |
| CDC91 | GCREF2013040 | 0.0078 | 0.125 | 4 | 0.125 | 16 | 2 | 1 |
| CDC92 | GCREF2013041 | 0.0039 | 0.125 | 0.125 | 0.031 | 16 | 2 | 1 |
| CDC93 | GCREF2013042 | 0.0039 | 0.125 | 0.125 | 0.016 | 16 | 1 | 1 |
| CDC94 | GCREF2013043 | 0.0078 | 2 | 0.125 | 0.008 | <0.016 | 0.25 | 0.5 |
| CDC95 | GCREF2013045 | 0.0078 | 0.125 | 0.25 | 0.031 | 16 | 2 | 1 |
| CDC96 | GCREF2013046 | 0.002 | 0.125 | 0.125 | 0.016 | 16 | 1 | 0.5 |
| CDC97 | GCREF2013047 | 0.0039 | 0.125 | 0.125 | 0.031 | 16 | 1 | 1 |
| CDC98 | GCREF2013048 | 0.0078 | 2 | 0.125 | 0.008 | <0.016 | 0.25 | 0.5 |
| CDC99 | GCREF2013012 | 0.0078 | 4 | 0.125 | 0.008 | <0.016 | 0.25 | 0.5 |
| CDC100 | GCREF2013015 | 0.0039 | 0.125 | 0.25 | 0.031 | 16 | 1 | 1 |

## REFERENCES

1. Guvenc, F., Kaul, R. & Gray-Owen, S. D. Intimate Relations: Molecular and Immunologic Interactions Between Neisseria gonorrhoeae and HIV-1. Front Microbiol 11, 1299, doi:10.3389/fmicb.2020.01299 (2020).

2. Prevention, C.-C. f. D. C. a. 2018 STD Surveillance Report Gonorrhea. (Division of STD Prevention, National Center for HIV/AIDS, Viral Hepatitis, STD, and TB Prevention, Centers for Disease Control and Prevention, 2019).

3. Unemo, M. Current and future antimicrobial treatment of gonorrhoea - the rapidly evolving Neisseria gonorrhoeae continues to challenge. BMC Infect Dis 15, 364, doi:10.1186/s12879-015-1029-2 (2015).

4. Lewis, D. A. New treatment options for Neisseria gonorrhoeae in the era of emerging antimicrobial resistance. Sex Health 16, 449–456, doi:10.1071/SH19034 (2019).

5. Unemo, M., Golparian, D. & Eyre, D. W. Antimicrobial Resistance in Neisseria gonorrhoeae and Treatment of Gonorrhea. Methods Mol Biol 1997, 37–58, doi:10.1007/978-1-4939-9496-0_3 (2019).

6. St Cyr, S. et al. Update to CDC’s Treatment Guidelines for Gonococcal Infection, 2020. MMWR Morb Mortal Wkly Rep 69, 1911–1916, doi:10.15585/mmwr.mm6950a6 (2020).

7. Unemo, M. & Shafer, W. M. Antimicrobial resistance in Neisseria gonorrhoeae in the 21st century: past, evolution, and future. Clin Microbiol Rev 27, 587–613, doi:10.1128/CMR.00010-14 (2014).

8. Unemo, M., Del Rio, C. & Shafer, W. M. Antimicrobial Resistance Expressed by Neisseria gonorrhoeae: A Major Global Public Health Problem in the 21st Century. Microbiol Spectr 4, doi:10.1128/microbiolspec.EI10-0009-2015 (2016).

9. (WHO)., W. H. O. Global health sector strategy on sexually transmitted infections 2016–2021: Towards ending STIs. Geneva: WHO; http://www.who.int/reproductivehealth/publications/rtis/ghss-stis/en/. (2016).

10. Drlica, K. & Zhao, X. DNA gyrase, topoisomerase IV, and the 4-quinolones. Microbiol Mol Biol Rev 61, 377–392 (1997).

11. Jacobsson, S. et al. High in vitro activity of the novel spiropyrimidinetrione AZD0914, a DNA gyrase inhibitor, against multidrug-resistant Neisseria gonorrhoeae isolates suggests a new effective option for oral treatment of gonorrhea. Antimicrob Agents Chemother 58, 5585–5588, doi:10.1128/AAC.03090-14 (2014).

12. Bradford, P. A., Miller, A. A., O’Donnell, J. & Mueller, J. P. Zoliflodacin: An Oral Spiropyrimidinetrione Antibiotic for the Treatment of Neisseria gonorrheae, Including Multi-Drug-Resistant Isolates. ACS Infect Dis 6, 1332–1345, doi:10.1021/acsinfecdis.0c00021 (2020).

13. Flamm, R. K., Farrell, D. J., Rhomberg, P. R., Scangarella-Oman, N. E. & Sader, H. S. Gepotidacin (GSK2140944) In Vitro Activity against Gram-Positive and Gram-Negative Bacteria. Antimicrob Agents Chemother 61, doi:10.1128/AAC.00468-17 (2017).

14. Biedenbach, D. J. et al. In Vitro Activity of Gepotidacin, a Novel Triazaacenaphthylene Bacterial Topoisomerase Inhibitor, against a Broad Spectrum of Bacterial Pathogens. Antimicrob Agents Chemother 60, 1918–1923, doi:10.1128/AAC.02820-15 (2016).

15. Damiao Gouveia, A. C., Unemo, M. & Jensen, J. S. In vitro activity of zoliflodacin (ETX0914) against macrolide-resistant, fluoroquinolone-resistant and antimicrobial-susceptible Mycoplasma genitalium strains. J Antimicrob Chemother 73, 1291–1294, doi:10.1093/jac/dky022 (2018).

16. Tari, L. W. et al. Tricyclic GyrB/ParE (TriBE) inhibitors: a new class of broad- spectrum dual-targeting antibacterial agents. PLoS One 8, e84409, doi:10.1371/journal.pone.0084409 (2013).

17. Jerse, A. E. et al. Estradiol-Treated Female Mice as Surrogate Hosts for Neisseria gonorrhoeae Genital Tract Infections. Front Microbiol 2, 107, doi:10.3389/fmicb.2011.00107 (2011).

18. Ohnishi, M. et al. Is Neisseria gonorrhoeae initiating a future era of untreatable gonorrhea?: detailed characterization of the first strain with high-level resistance to ceftriaxone. Antimicrob Agents Chemother 55, 3538–3545, doi:10.1128/AAC.00325-11 (2011).

19. Tacconelli, E. et al. Discovery, research, and development of new antibiotics: the WHO priority list of antibiotic-resistant bacteria and tuberculosis. Lancet Infect Dis 18, 318–327, doi:10.1016/S1473-3099(17)30753-3 (2018).

20. Belland, R. J., Morrison, S. G., Ison, C. & Huang, W. M. Neisseria gonorrhoeae acquires mutations in analogous regions of gyrA and parC in fluoroquinolone- resistant isolates. Mol Microbiol 14, 371–380, doi:10.1111/j.1365-2958.1994.tb01297.x (1994).

21. Lindback, E., Rahman, M., Jalal, S. & Wretlind, B. Mutations in gyrA, gyrB, parC, and parE in quinolone-resistant strains of Neisseria gonorrhoeae. APMIS 110, 651–657, doi:10.1034/j.1600-0463.2002.1100909.x (2002).

22. Bisacchi, G. S. & Manchester, J. I. A New-Class Antibacterial-Almost. Lessons in Drug Discovery and Development: A Critical Analysis of More than 50 Years of Effort toward ATPase Inhibitors of DNA Gyrase and Topoisomerase IV. ACS Infect Dis 1, 4–41, doi:10.1021/id500013t (2015).

23. Oblak, M., Kotnik, M. & Solmajer, T. Discovery and development of ATPase inhibitors of DNA gyrase as antibacterial agents. Curr Med Chem 14, 2033–2047, doi:10.2174/092986707781368414 (2007).

24. Charifson, P. S. et al. Novel dual-targeting benzimidazole urea inhibitors of DNA gyrase and topoisomerase IV possessing potent antibacterial activity: intelligent design and evolution through the judicious use of structure-guided design and structure-activity relationships. J Med Chem 51, 5243–5263, doi:10.1021/jm800318d (2008).

25. Tomasic, T. et al. Discovery of 4,5,6,7-Tetrahydrobenzo[1,2-d]thiazoles as Novel DNA Gyrase Inhibitors Targeting the ATP-Binding Site. J Med Chem 58, 5501–5521, doi:10.1021/acs.jmedchem.5b00489 (2015).

26. Gjorgjieva, M. et al. Discovery of Benzothiazole Scaffold-Based DNA Gyrase B Inhibitors. J Med Chem 59, 8941–8954, doi:10.1021/acs.jmedchem.6b00864 (2016).

27. Basarab, G. S. et al. Responding to the challenge of untreatable gonorrhea: ETX0914, a first-in-class agent with a distinct mechanism-of-action against bacterial Type II topoisomerases. Sci Rep 5, 11827, doi:10.1038/srep11827 (2015).

28. Alm, R. A. et al. Characterization of the novel DNA gyrase inhibitor AZD0914: low resistance potential and lack of cross-resistance in Neisseria gonorrhoeae. Antimicrob Agents Chemother 59, 1478–1486, doi:10.1128/AAC.04456-14 (2015).

29. Miller, A. A., Traczewski, M. M., Huband, M. D., Bradford, P. A. & Mueller, J. P. Determination of MIC Quality Control Ranges for the Novel Gyrase Inhibitor Zoliflodacin. J Clin Microbiol 57, doi:10.1128/JCM.00567-19 (2019).

30. Bax, B. D. et al. Type IIA topoisomerase inhibition by a new class of antibacterial agents. Nature 466, 935–940, doi:10.1038/nature09197 (2010).

31. Jones, R. N., Fedler, K. A., Scangarella-Oman, N. E., Ross, J. E. & Flamm, R. K. Multicenter Investigation of Gepotidacin (GSK2140944) Agar Dilution Quality Control Determinations for Neisseria gonorrhoeae ATCC 49226. Antimicrob Agents Chemother 60, 4404–4406, doi:10.1128/AAC.00527-16 (2016).

32. Biedenbach, D. J. et al. In Vitro Activity of AZD0914, a Novel Bacterial DNA Gyrase/Topoisomerase IV Inhibitor, against Clinically Relevant Gram-Positive and Fastidious Gram-Negative Pathogens. Antimicrob Agents Chemother 59, 6053–6063, doi:10.1128/AAC.01016-15 (2015).

33. Prevention, C.-C. f. D. C. a. Antibiotic resistance threats in the United States, 2013. Atlanta, GA, CDC. 2013. http://www.cdc.gov/drugresistance/threat-report-2013/pdf/ar-threats-2013-508.pdf. (2013).

34. Jerse, A. E. Experimental gonococcal genital tract infection and opacity protein expression in estradiol-treated mice. Infection and immunity 67, 5699–5708 (1999).

35. Connolly, K. L. et al. Pharmacokinetic Data Are Predictive of In Vivo Efficacy for Cefixime and Ceftriaxone against Susceptible and Resistant Neisseria gonorrhoeae Strains in the Gonorrhea Mouse Model. Antimicrob Agents Chemother 63, doi:10.1128/AAC.01644-18 (2019).

36. Song, W. et al. Local and humoral immune responses against primary and repeat Neisseria gonorrhoeae genital tract infections of 17beta-estradiol-treated mice. Vaccine 26, 5741–5751, doi:10.1016/j.vaccine.2008.08.020 (2008).

37. Ochsner, U. A. et al. Inhibitory effect of REP3123 on toxin and spore formation in Clostridium difficile, and in vivo efficacy in a hamster gastrointestinal infection model. J Antimicrob Chemother 63, 964–971, doi:10.1093/jac/dkp042 (2009).

